# Behavioural responses to acute warming precede critical shifts in the cellular and physiological thermal stress responses in fish

**DOI:** 10.1101/2024.01.29.577477

**Authors:** Travis C. Durhack, Matt J. Thorstensen, Theresa E. Mackey, Mélanie Aminot, Michael J. Lawrence, Céline Audet, Eva C. Enders, Ken M. Jeffries

## Abstract

From a conservation perspective, it is important to identify when sub-lethal temperatures begin to adversely impact an organism. However, it is unclear whether, during acute exposures, these cellular thresholds occur at similar temperatures to other physiological or behavioural changes. To test this, we estimated temperature preference (15.1 ± 1.1 °C) using a shuttle box, thermal optima for aerobic scope (10–15 °C) using respirometry, agitation temperature (22.0 ± 1.4 °C) as the point where a fish exhibits a behavioural avoidance response and the *CT_max_* (28.2 ± 0.4 °C) as the upper thermal limit for 1 yr old Brook Trout (*Salvelinus fontinalis*) acclimated to 10 °C. We then acutely exposed a different subset of fish to these temperatures and sampled tissues when they reached the target temperature or after 60 min of recovery at 10 °C. We used qPCR to estimate mRNA transcript levels of genes associated with heat shock proteins, oxidative stress, apoptosis, and inducible transcription factors. A major shift in the transcriptome response occurred near the agitation temperature, which may identify a link between the cellular stress response and the behavioural avoidance response.

## Introduction

Many riverine systems are experiencing increasing water temperatures due to climate change and habitat modifications (Bowerman et al., 2021; Mauger et al., 2017; Morash et al., 2020; Westley, 2020). Changes in temperature affect freshwater fishes, with increased summer temperatures in river systems leading to increased mortality in fishes, including migratory populations (Bowerman et al., 2021; Mauger et al., 2017; Morash et al., 2020; Westley, 2020) and juvenile resident populations that are unable to escape unfavourable temperatures (Cairns et al., 2005; Morash et al., 2020; Petty et al., 2012; Rodnick et al., 2008; White & Wagner, 2021). Understanding a species’ or population’s temperature preference and upper thermal thresholds is important for determining how increasing water temperatures will affect wild fish populations.

There are several common methods for estimating the thermal tolerance limits in fishes that assess endpoints at different levels of biological organization. The critical thermal maximum (*CT_max_*) is a commonly used non-lethal estimate of its acute upper thermal limits for a species or population, while the upper incipient lethal temperature (UILT) estimates a longer term upper thermal threshold (Beitinger et al., 2000). Acclimation to differing temperature regimes can affect the thermal tolerance of fishes, with previous studies showing differences in *CT_max_* within the same population of fish acclimated to different temperatures (Kelly et al., 2014; Morrison et al., 2020). A species’ preferred temperature often coincides with their optimal growth and metabolism (Macnaughton et al., 2021; Schulte et al., 2011), and these physiological parameters can often be found to correspond with increased activities and behaviours, such as feeding, reproduction, or avoidance responses (Crawshaw, 1977; Killen et al., 2013). Temperatures around a species’ preferred temperature generally provide the greatest opportunity for energetically expensive activities (e.g., growth and reproduction), while temperatures towards the upper and lower ends of a species’ thermal range tend to reduce or prevent them (Crawshaw, 1977, 1984; Desforges et al., 2023). Research assessing the *CT_max_*of fishes has shown an activation of a behavioural response at temperatures leading up to *CT_max_*, which has been described as the agitation temperature (McDonnell & Chapman, 2015). Additionally, the *CT_max_*-agitation window, which represents the difference between the temperatures for these endpoints can be used to assess how early a species’ avoidance response is activated before critical temperatures are reached (Wells et al., 2016). Studies that assess thermal endpoints can provide a general understanding of when fish display a behavioural reaction to a thermal threshold, however, such studies do not provide insight into sub-organismal responses of the fish. Despite the potential utility of identifying behavioural avoidance responses to thermal stress in fishes, relatively little is known about the physiological mechanisms that coincide with the agitation temperature in fishes (Bouyoucos et al. 2023).

Measuring how temperature affects performance traits can be accomplished using thermal performance curves (TPC). Performance traits, defined as biological processes that occur over a period of time (e.g., growth, reproduction, metabolic rate; Schulte et al., 2011), often peak at an optimal temperature and decrease as the temperature moves further away from this optimum. Estimates of the amount of energy available to an organism, both for basic biological functions (e.g., digestion and growth) or energy demanding life processes (e.g., reproduction), can be drawn from the determination of aerobic scope (AS) (Fry, 1947). Metabolic rates of fishes are often estimated using oxygen consumption over a set period of time, with standard metabolic rate (SMR) representing the lowest oxygen consumption or minimum energy requirements to sustain physiological function (i.e., homeostasis; Beamish & Mookherjii, 1964; Brett & Groves, 1979; Fry, 1971), maximum metabolic rate (MMR) being the highest rate of oxygen consumption, often measured following exhaustive exercise (Brett & Groves, 1979; Treberg et al., 2016), and AS signifying the difference between SMR and MMR (Fry, 1947). Requirements and availability of energy for fish changes with water temperature, and an optimal temperature for aerobic performance is often considered to be where AS peaks. It is important to note that a defined AS peak is not always present in fishes with some species able to maintain AS over a broad range of temperatures (e.g., Brook Trout (*Salvelinus fontinalis*); Durhack et al., 2021), possibly right up to lethal temperatures (Clark et al., 2013; Munday et al., 2009). However, at temperatures above peak AS, cardiac arrhythmia, and increased reliance on anaerobic metabolism have been shown to occur in salmonid species (Anttila et al., 2013) highlighting the importance of investigating responses to temperature across different levels of biological organization. Combining whole-body metabolic, behavioural, and cellular responses to changes in water temperature provides a full picture of the integrated organismal response to different temperatures, from their preferred temperature to thermal extremes.

When an organism experiences changes in their thermal environment, the activation of the heat shock response (HSR) via up- or downregulation of heat shock proteins (HSPs) occurs (Tomanek, 2010). However, acclimation to a temperature within the thermal limits of a fish can change when the HSR is induced. Within the thermal distribution of a species, moderate temperature increases may be stimulatory, with a potential benefit to the organism (Jeffries et al., 2016, 2018; Schreck, 2010). As temperatures increase there will be a sub-lethal threshold where a thermal stress response is activated, which may lead to a reduction in organismal fitness (Buckley et al., 2006; Komoroske et al., 2015; Logan & Somero, 2011; Schulte, 2014). At extreme temperatures approaching *CT_max_* an individual will start exhibiting a HSR (Jeffries et al., 2018). Exposure to these extreme temperatures is where cellular level damage may take place leading to apoptosis and there is an up- and downregulation of genes associated with cellular survival mechanisms (Komoroske et al., 2015; Logan & Somero, 2011). Consequently, recovery from exposure to extreme temperatures may be delayed or no longer possible. Survival at acute exposures to temperatures above the onset of a thermal stress response is possible, however, prolonged exposure to temperatures on the high end of a species’ TPC may have impacts on fitness.

The aim of this study was to assess whether the behavioural and whole-organism responses relate to shifts in the cellular response to acute temperature increases. Brook Trout are a eurythermal species that are widely distributed throughout North America. Upper thermal tolerance and temperature preference studies have been previously conducted on other populations of this species and temperature preference, peak AS, UILT, and *CT_max_* have been estimated to be 15.9 ± 1.4 °C, 15 °C, 25.1 ± 0.2 °C, and 29.9 ± 0.6 °C respectively (values from: Durhack et al., 2021; Hasnain, 2012; Macnaughton et al., 2021; Morrison et al., 2020; reviewed by Smith & Ridgway, 2019). We first conducted intermittent respirometry, shuttle box, and *CT_max_*experiments to identify key behavioural and physiological endpoints commonly used to study temperatures impacts on fishes. Following a *CT_max_*experimental protocol, we acutely exposed a different subset of fish from the same cohort to different target temperatures ranging from the preferred temperature to the *CT_max_*. Gill, liver, and blood samples were then taken from fish to quantify the changes in mRNA transcript levels and plasma indices. The combination of experiments to assess responses to changes in temperature at a whole body, cellular, and behavioural level tested the hypothesis that there is a sub-lethal threshold at a temperature approaching the acute upper thermal limit (i.e., *CT_max_*) characterized by both behavioural and cellular responses. Defining critical physiological thresholds could be used as an upper temperature limit for designating suitable habitat for a species in the wild.

## Methods

### Animal care and holding

Young-of-the-year Brook Trout were obtained from the Whiteshell Fish Hatchery in southeastern Manitoba, Canada. The strain originated from Lake Nipigon, Ontario, Canada and was brought to the hatchery in the summer of 2019 (P1), at which point a batch was obtained and held at Fisheries and Oceans Canada’s Freshwater Institute fish holding facility in 600 L tanks that were maintained on a flow-through of de-chlorinated City of Winnipeg tap water and with independent aeration. Fish were maintained under a 12-12 photoperiod (65 min of dawn/dusk, full light starting at 07:05, and full dark at 19:05) at 10 °C ± 1 °C. Fish were fed daily on a diet of commercial food pellets (EWOS Pacific: Complete Fish Feed for Salmonids, Cargill, Minneapolis, MN, USA). All procedures were conducted under Animal Use Protocols approved by the Fisheries and Oceans Canada Freshwater Institute Animal Care Committee (FWI-ACC AUP-2020-05 & FWI-ACC AUP-2020-07) under the standards set by the Canadian Council for Animal Care.

### Experimental Treatments

#### Temperature preference experiments

A total of *n* = 15 fish were tested for temperature preference using the Shuttle Box system (Loligo^®^ Systems, Viborg, Denmark). Fish were haphazardly selected from the general population tank after feeding and transferred into the Shuttle Box for 48 h. Fish were allowed to acclimate to the shuttle box tank for ∼15 min before the system was turned on and movement tracking started. The temperature in the system started at 10 °C when the fish was put in the tank and was maintained between a maximum of 25 °C and a minimum of 5 °C for the safety of the fish, as these temperatures are within the temperature tolerance of Brook Trout (reviewed by Smith & Ridgway, 2019). Fish were allowed to freely swim back and forth between the “heating” and “cooling” chambers on either side of the tank, and temperature difference between the “heating” and “cooling” chambers was kept within 2 °C of each other, as well as having a maximum temperature change of 3 °C per hour to avoid heat stressing the fish. Room lighting was kept on a 12-12 cycle, the same as in the general population tank. During low light/overnight times, infrared-light was used to illuminate the tank from below to allow the camera to track the fish. Infrared lights were on a timer to turn on at 18:30 and turn off at 07:30, which coincided with dawn and dusk periods of the room lighting cycle. It was noted that for ∼15 min at these times the camera was unable to track the fish, however, tracking was successful during the rest of the day/night and no significant changes in temperature occurred during these brief windows. Fish movement was monitored by an overhead camera and water temperature was recorded using a TMP-REG instrument (Loligo^®^ Systems, Viborg, Denmark) constantly for 48 h by the Shuttlesoft software (Loligo^®^ Systems, Viborg, Denmark). The software then automatically calculated a preferred temperature based on the water temperature that the fish sought out. The first 24 h that the fish was in the Shuttle Box was considered an acclimation period, and the data from this time was excluded from final analysis of temperature preference.

#### Intermittent Respirometry experiments

From December 14, 2020 – March 17, 2021, we conducted intermittent respirometry experiments on *n* = 65 fish to estimate metabolic rates (SMR, MMR and AS). Groups of *n* = 15 fish were acclimated for a minimum of three weeks at 5, 10, 15, 20 and 23 ± 0.1 °C. Intermittent respirometry was conducted using AutoResp software to monitor and record oxygen and temperature levels via Witrox4 instruments and control water pumps via a DAQ-M instrument (Loligo^®^ Systems, Viborg, Denmark). Fish were fasted for a minimum of 24 h prior to experimentation. Following fasting, one fish at a time was haphazardly netted from the general population tanks and first tested for MMR using an exhaustive chase protocol. The exhaustive chase protocol entailed fish being coaxed to swim against a constant current of water until they were deemed exhausted. Exhaustion was determined to be once a fish no longer responded to a gentle pinch of the caudal fin (Durhack et al., 2021; Reidy et al., 1995). Fish were then immediately transferred to acrylic respirometry chambers (volume 679 mL; Loligo^®^ Systems, Viborg, Denmark) where three measurements were recorded (measurement cycle = Measure – 180 s, Flush – 300 s, Wait – 40 s) to estimate MMR. The flush and wait periods were skipped for the first measurement to ensure no recovery period was allowed for the fish before measurements began. Following MMR estimates, □O_2_ measurements were recorded for 24 h to be used for SMR estimates. Following SMR, fish were sacrificed as described below, measured for fork and total length and sex determination.

#### CT_max_ experiments

Treatments of *n* = 16 fish were used in this study (total of *n* = 208). Six treatments (handling, acclimation, agitation, *CT_max_*, B1 and B2) were separated into two time points: T^0^ – where fish were sampled immediately once either a certain temperature or behaviour was reached, or T^1^ – where the fish were given a 1 h recovery period in a recovery bath held at 10 □ once the temperature or behaviour was reached (Figure 1). A group of *n* = 16 fish was also used as a control group to assess baseline stress levels. This group will here forth be termed the “baseline” treatment. Fish were tested in groups of 4 in one of two 200 L green sampling tanks, each in an individual acrylic chamber (volume – 1380 mL) with netting over the ends to contain and track the fish (Figure 2). For treatments other than the baseline group, fish were haphazardly netted from the general population tank and placed in their sampling chambers to fast and acclimate overnight prior to experimental trials (Approx. 18 hours). Temperature of the tank was monitored using a Witrox 4 instrument and AutoResp software (Loligo® Systems, Viborg, Denmark). Both sampling tanks had air stones to maintain dissolved oxygen levels and two 5 L·min^-1^ pumps (Eheim, Deizisau, Germany) to ensure homogenous temperature throughout the tank. For treatments where heating was needed, a rate of approximately 0.3 °C min^-1^ was used with four 300 W heating elements (Finnex TH-0300S titanium heaters, Finnex, Chicago, USA). Fish were removed from the treatment and either immediately sacrificed (T^0^) or moved to the recovery bath (T^1^) when they either reached the specified temperature or exhibited the specified behaviour. For treatments that ended with a specified temperature, fish were removed two at a time, with one fish designated T^0^ for sampling and one fish designated for T^1^ and placed into the recovery bath. The remaining two fish were held at stable temperature (± 0.2 °C) for 2 min to allow time for tissue sampling from the first fish. In treatments with a behavioral endpoint, fish were removed once the behaviour was noted and sampled in the same order as above. Behavioral endpoint fish were not necessarily spaced out as well as temperature endpoint fish since sampling depended on when the individual fish reached the endpoint (Figure 1). All fish were sacrificed in Syncaine (MS-222; concentration: 450 mg L^-1^ buffered with 900 mg L^-1^ sodium bicarbonate; Syndel Canada, Nanaimo, BC, Canada) for 3 min followed by cranial percussion. Oxygen saturation was measured at the beginning, middle, and end of each trial and never fell below 98%.

**Figure 1.**
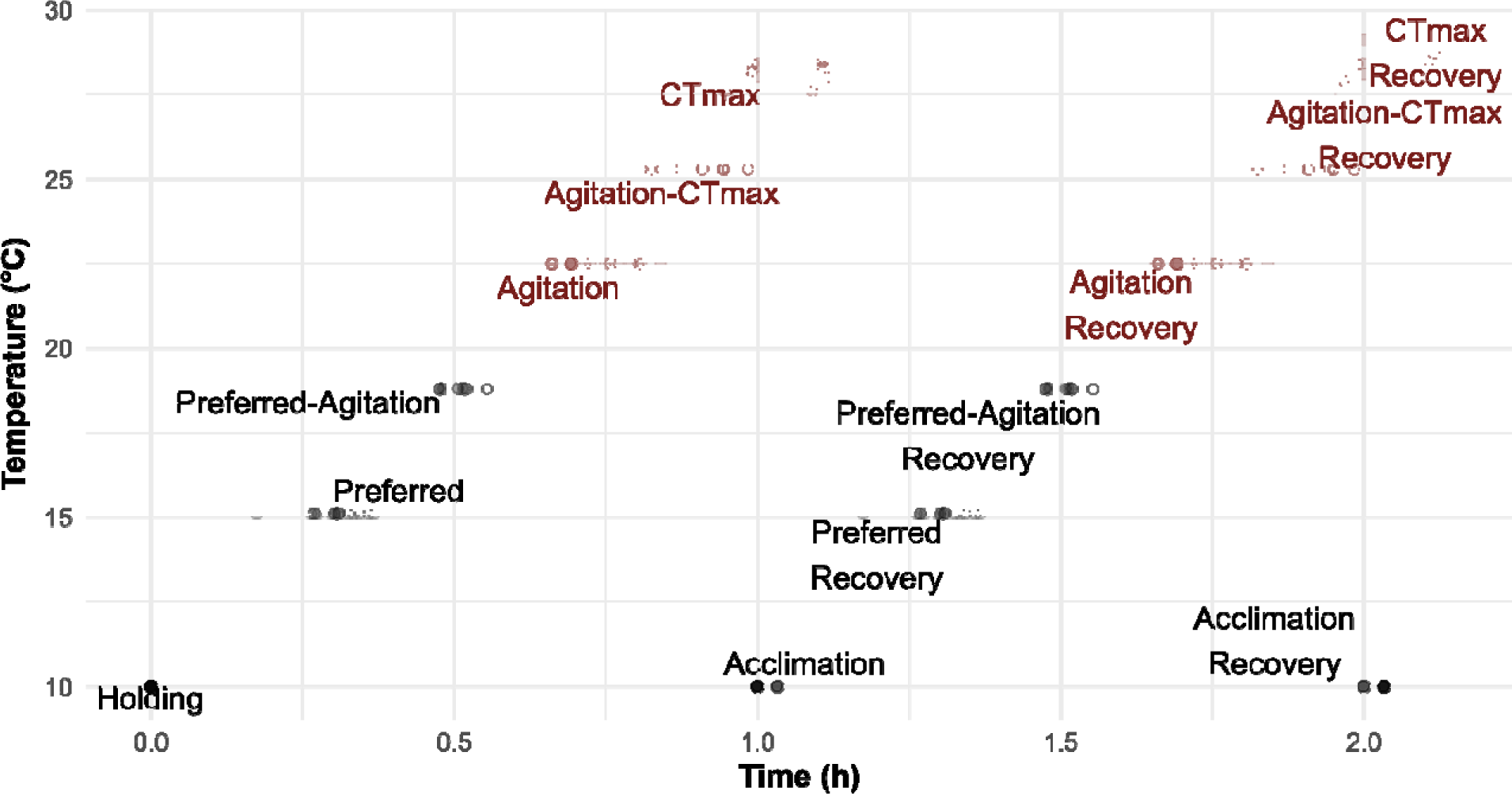
Experimental design of treatments and timing for gene expression tissue sampling. Fish were sampled for white muscle, liver, and gill. Each treatment was sampled either directly following heating to the treatment temperature or following a h recovery period in a 10 °C recovery tank.

**Figure 2.**
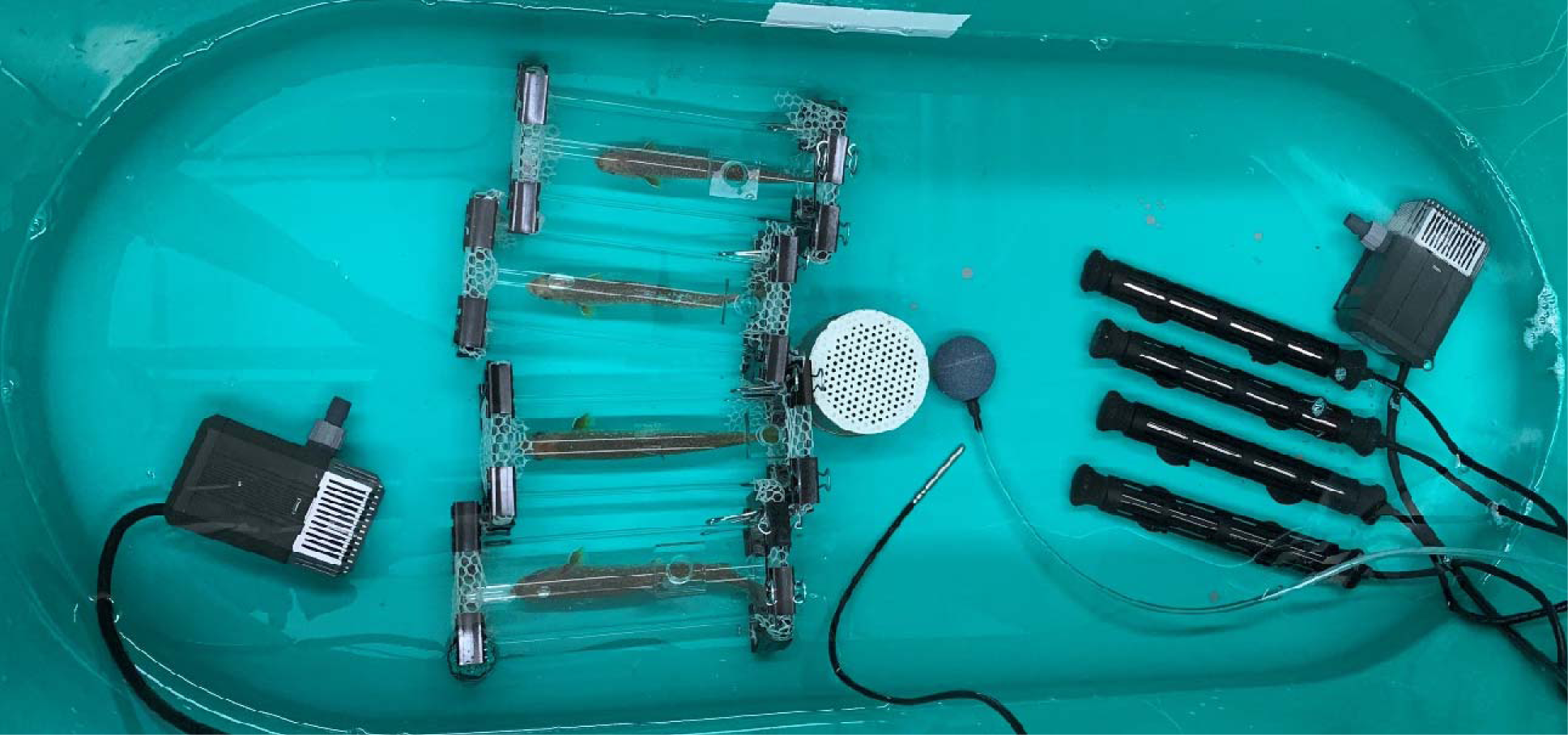
Example of *CT_max_*experimental setup. Treatments of *n* = 16 fish were used in this study (total of *n* = 208). Six treatments (handling, acclimation, agitation, *CT_max_*, B1 and B2) were separated into two time points: T^0^ – where fish were sampled immediately once either a certain temperature or behaviour was reached, or T^1^ – where the fish were given a 1 h recovery period in a recovery bath held at 10 □ once the temperature or behaviour was reached (Figure 1). A group of *n* = 16 fish was also used as a control group to assess baseline stress levels. Fish were tested in groups of 4 in one of two 200 L green sampling tanks, each in an individual acrylic chamber (volume – 1380 mL) with netting over the ends to contain and track the fish. Each tank also contained an air stone to maintain dissolved oxygen levels above 90%, a temperature probe to monitor heating rates, four 300 watt titanium heaters and two 5L/min pumps for water movement to ensure even heating throughout the tank.

#### Description of CT_max_ treatment groups

The control groups are described first. The fish for the baseline treatment were haphazardly netted out of the general population tanks and immediately euthanized for sampling as described above. The baseline treatment fish were used to assess baseline stress levels before fish are introduced to the *CT_max_* chambers. Another control group, termed the “handling” control group, was treated the same as all experimental treatments, but instead of the water being heated, the tank was kept at a temperature of 10 °C for one hour (∼ length of time for the *CT_max_* treatment). T^0^ fish were sampled to assess stress levels from experimental housing without temperature effects, while T^1^ fish represent residual stress levels from handling in fish following 1 h of recovery.

The preferred temperature treatment for this population of fish was established with the shuttle box experiments described above. The temperature preference experiments determined the average preferred temperature to be 15.1 ± 1.13 °C. As such, 15.1 °C was used as the target temperature for this treatment.

For the agitation treatment, we defined the agitation temperature as the temperature at which sustained (>5 s) ‘curling’ and ‘bursting’ behaviours were displayed by the fish (Video 1). The curling behaviour was defined as the fish’s body curved in a C-shape and was a result of attempting to turn around in the sampling chambers to look for an escape from the water temperature. Bursting behaviour was defined as bursts of energetic swimming by the fish and was often observed as sustained pushing against the netting enclosing the ends of the chamber. Again, this was likely an attempt to escape increasing water temperatures. We chose these two behaviours because we sought to study a flight response possibly reflected in physiological changes in a stressed fish, as opposed to a less extreme preference response with less drastic movements (McDonnell & Chapman, 2015). Preliminary assessment of 16 fish from *CT_max_* trials led us to define the average agitation temperature for this treatment as 22.5 °C. The average agitation temperature was used to define temperatures for treatments B1 and B2.

The B1 treatment was defined as the midpoint between the preferred temperature treatment – 15.1 °C – and the mean agitation temperature treatment – 22.5 °C. As such, fish in the B1 treatment were sampled at 18.8 °C.

As with the B1 treatment, the B2 treatment was defined as a group between the mean agitation temperature and the average *CT_max_* temperature. The average *CT_max_* temperature was estimated from the first *n* = 16 fish to undergo a CT_max_ trial and was found to be 28.1 °C. The B2 treatment temperature was thus set at 25.3 °C, halfway between the mean agitation temperature – 22.5 °C – and the average *CT_max_* – 28.1 °C.

Similar to the agitation temperature treatment, we sampled fish in the *CT_max_* treatment at their individual *CT_max_*thresholds. Fish were deemed to have reached *CT_max_* once they were unable to maintain equilibrium and ceased attempts to right themselves. In each trial of four, the first fish to reach *CT_max_* was sampled at T^0^, the second went to recovery for T^1^, the third sampled for T^0^, and the fourth to recovery for T^1^ so that neither the T^0^ or T^1^sampling groups would be biased for fish with low or high *CT_max_* values.

#### Tissue Sampling

Following euthanasia, fish length and mass were recorded. Blood was then collected using ammonium-heparinized capillary tubes (Fisherbrand^®^, Fisher Scientific, Pittsburgh, Pennsylvania, USA) following severing of the caudal fin. Gill tissue from the ventral side of the second gill arch, cut to the base of the gill filaments was taken for each fish. Liver was excised, weighed, and the distal end of the lobe was taken. Each tissue was immediately stored in RNA*later*^TM^ (Invitrogen™, Carlsbad, California, USA) kept on ice, and stored at 4 °C overnight prior to storage at −80 °C following RNA*later*^TM^ best practices (Life Technologies, 2011). Following this, the sex of the animal was determined visually (male, female, or indeterminant).

#### Quantitative PCR

Total RNA was extracted from the gill and liver tissues using a Qiagen RNeasy Plus Mini Kit (Qiagen, Toronto, ON, CA) following manufacturer’s protocols. The RNA samples were checked for purity (A260/A280, A260/A230) and concentration using a NanoDrop One Spectrophotometer (Thermo Fisher Scientific, Wilmington, DE, USA). The integrity of the RNA was assessed by electrophoresis on a 1% agarose gel. 1 µg of total RNA was reverse transcribed into cDNA using the Invitrogen^TM^ SuperScript^TM^ IV First-Strand Synthesis System (Thermo Fisher Scientific, Waltham, MA, USA) following the manufacturer’s protocols.

Forward and reverse quantitative PCR (qPCR) primers and probes (*cat, cirbpa, ef1a, fos, gpx1a, hsf1, hsp70a, hsp90aa, hsp90ba, ier2, jun, junb, jund, mycb, rpl7, rpl13a, rps9, sod1, sod2*; Table 1) were designed in Geneious Prime software version 2021.2.2 (Biomatters Ltd, Auckland, New Zealand) based off multiple salmonid species using sequences from GenBank® and PhyloFish (Sayers et al., 2019; Sutherland et al., 2019). The remaining primers and probes (*casp9, cs, ldh, hsp90ab1*; Supplementary Table 1) were designed by Jourdain-Bonneau et al. (2023) using Primer Express software version 3.0 (Applied Biosystems, ThermoFisher Scientific, Wilmington, DE, USA).

**Table 1.** Mass specific metabolic rate estimates ± standard error (mg O^2^ kg^-1^ h^-1^) for juvenile Brook Trout assessed using intermittent respirometry.

| Temperaure (°C) | Standard Metabolic Rate (mg O<sup>2</sup>·kg<sup>-1</sup>·h<sup>-1</sup>) | Maximum Metabolic Rate (mg O<sup>2</sup>·kg<sup>-1</sup>·h<sup>-1</sup>) | Aerobic Scope (mg O<sup>2</sup>·kg<sup>-1</sup>·h<sup>-1</sup>) |
| --- | --- | --- | --- |
| 5 | 31.02 ± 5.03 | 358.90 ± 17.00 | 300.63 ± 17.37 |
| 10 | 38.27 ± 2.51 | 477.71 ± 17.19 | 396.30 ± 18.00 |
| 15 | 72.56 ± 3.92 | 498.66 ± 13.01 | 387.54 ± 12.76 |
| 20 | 105.40 ± 10.78 | 392.64 ± 19.88 | 242.45 ± 14.28 |
| 23 | 174.23 ± 25.64 | 325.50 ± 35.14 | 138.93 ± 42.70 |

Primers and probes were designed for 18 target genes (Supplementary Table 1) that represented transcriptomic responses consistent with high temperature (*cold-inducible RNA-binding protein* (*cirbp*), *heat shock transcription factor-1* (*hsf1*), *heat shock protein 70-alpha* (*hsp70a*)*, cytosolic heat shock protein 90-alpha* (*hsp90aa*), *heat shock protein 90-beta-1* (*hsp90ab1*), *heat shock protein 90-beta-alpha* (*hsp90ba*)), cell cycle and transcription (*caspase-9* (*casp9*), *catalase* (*cat*), *citrate synthase* (*cs*), *protein c-fos* (*fos*), *immediate-early response gene 2* (*ier2*), *transcription factor ap-1* (*jun*), *transcription factor jun-b* (*junb*), *transcription factor jun-d* (*jund*), *transcriptional regulator myc-2* (*mycb*)), and general cellular function (*glutathione peroxidase 1a* (*gpx1a*), *lactate dehydrogenase* (*ldh*)). Primers were designed for four reference genes including *elongation factor 1-alpha* (*ef1a*), 60*s ribosomal protein L13a* and *L7* (*rpl13a* and *rpl7*), and *40s ribosomal protein S9* (*rps9*) (Supplementary Table 1). Primer and probe efficiencies were tested by generating standard curves using cDNA synthesized from the RNA pooled from 7 individuals from the treatment groups. Each 12 µL qPCR reaction consisted of 1 µL of a 1:10 dilution of cDNA, 500 nM forward and reverse primer, 6 µL of Applied Biosystems^TM^ Taqman FAST Advanced Master Mix (Thermo Fisher Scientific, Waltham, MA, USA) 0.3 µL of 10 µM Taqman probe, and 0.34 µL RNase-free water. The qPCR reactions were run on a QuantStudio 5 Real-Time PCR System (Thermo Fisher Scientific, Life Technologies Corporation, Carlsbad, California USA) in 384 well plates.

## Analysis

Temperature preference for each fish was determined by taking the final value of the recorded temperature preference values from the Shuttlesoft software (Loligo^®^ Systems, Viborg, Denmark). Overall temperature preference was determined by taking the mean ± S.D. of all fish tested. We tested for differences in temperature preference of juvenile Brook Trout between daytime and nighttime hours, as previous studies had seen differences in temperature preference between daytime and nighttime using the same system for another salmonid (Westslope Cutthroat Trout, *Oncorhynchus clarkii lewisi*; Macnaughton et al., 2018). No differences were found, as such we included all 24 h of data to estimate our final temperature preference.

### Metabolic Rate

Variation in fish body mass ranging from 27.8–171.1 g – was observed within the fish used in the respirometry experiment. To account for the range of fish mass across treatments and the known effect of mass on metabolic rate, whole body SMR, MMR, and AS estimates were mass corrected to the mean mass of all fish in the study (79.08 g) using multivariate polynomial predictive equations (similar to Durhack et al., 2021; Poletto et al., 2017). Sex was also determined for these fish to allow for assessment of metabolic differences between male (*n* = 43) and female fish (*n* = 32), respectively, and all fish were deemed to be immature. The inter-individual variability within our sampled fish allowed for testing of possible differences and interactions between the variables mass, sex and time to exhaustion and their possible effects on metabolic rate estimates. Tukey’s honest significant difference test (Tukey HSD) was used for post-hoc testing on any significant variables found to identify differences in temperature, mass, time to exhaustion and sex within and across treatments, with a P < 0.05 deemed significant. Mass-corrected data are presented in mass specific values (mg O^2^⋅kg^-1^ h^-1^) for comparison to previous studies. Statistical analysis was conducted using R and R Studio (RStudio version 1.3.1056, Posit Team, 2020; R version 4.0.2, R Core Team, 2020) with the following packages ‘car’ (Fox & S., 2019), ‘caret’ (Kuhn, 2008), ‘dplyr’ (Wickham et al., 2020), ‘fishMO2’ (Chabot et al., 2016), ‘MASS’ (Venables & Ripley, 2002), ‘multcomp’ (Hothorn et al., 2008), ‘MuMin’ (Barton, 2020), ‘plotrix’ (J, 2006), and ‘tidyverse’ (Wickham et al., 2019).

### CTmax

*CT_max_* and agitation temperatures were determined by taking the mean ± standard deviation values of all treatments tested at least up that temperature. Estimation of *CT_max_* was only taken from fish fully heated to *CT_max_*, which included our original 16 fish used to define the other treatment temperatures and the *CT_max_* treatment. As such, *n* = 32 fish were used to estimate *CT_max_*. Agitation temperature was assessed using all fish from the *CT_max_*, Agitation and B_2_ treatments, resulting in *n* = 96 fish. The agitation-*CT_max_* window was determined as the difference between the overall means of *CT_max_* and agitation temperatures.

### Plasma Lactate

A linear model was used to assess the extent to which plasma lactate concentration in nmol·µL^-1^ changed across experimental treatments and between the T^0^ and T^1^ groups. The model consisted of lactate concentration dependent on time in minutes from the start of experimental manipulations to blood draw (sampling time), Fulton’s condition factor, sex, and the interaction of experimental treatment and recovery time. Model fit was assessed with the *check_model()* function from the R package performance v0.8.0 (Lüdecke et al., 2021). Effect size statistics were calculated with the *anova_stats()* function of the R package sjstats v0.18.1 (Lüdecke, 2018) on a type II ANOVA performed on the linear model with the R package car v3.0-12 (Fox & Weisberg, 2019). 95% confidence intervals given the interaction between experimental treatment and recovery time were calculated with predicted marginal effects for each experimental treatment except baseline with sjPlot v2.8.10 (Lüdecke, 2021). Fish in the baseline treatment were omitted because the interaction was rank deficient since no T^1^ fish were sampled for that group.

### qPCR Data

The R package MCMC.qpcr v1.2.4 with MCMCglmm v2.33 was used as a Bayesian approach designed for qPCR data (Hadfield, 2010; Matz et al., 2013). Four reference genes were used in all models: *rpl13a*, *rps9*, *rpl17*, and *ef1a*. Stability of the reference genes across treatments was confirmed visually prior to including them in the analyses. Separated liver and gill models were used because 20 genes total were analyzed in liver and 22 total in gill. Transcript abundance of *jund*, *junb*, and *mycb*, were analyzed in the gill but not liver, while *hsp90aa* was analyzed in the liver but not gill. Within each tissue, treatment and recovery conditions were treated as interacting variables. The baseline treatment was omitted because the interaction was rank deficient, as no fish from baseline were in the T^1^ recovery group. The T^0^ and the acclimation groups were used as references against which other treatments were compared.

In all models, sex, sampling time, and Fulton’s condition factor were included as fixed effects that may alter transcript abundance estimates or gene expression during sampling. Each of the models was run with Markov Chain Monte Carlo settings of 110,000 iterations run total, a burn-in period of 10,000 iterations discarded, and a thinning interval of parameters sampled once every 100 iterations. Model convergence was assessed by observing trace plots and overall diagnostic plots of residuals and predicted values, variance of the residuals and predicted values (as a test of homoscedasticity), and normality of residuals. Differences in transcript abundance between treatments were assessed with 95% credible intervals. The package tidybayes v3.0.2 was used to visualize posterior estimates of transcript abundance (Kay, 2021). Separate linear models were used to assess the possibility that changes in mRNA abundance may reflect sampling time (Supplemental Methods).

## Results

### Intermittent Respirometry

Both AS and MMR peaked between 10–15 □ (Figure 3-A & -B), with 10 and 15 □ values being significantly different from all other treatment temperatures. However, no differences were found between temperature treatments for either AS or MMR at 10 and 15 □ (Tukey, *p* = 1.0, *p* = 0.95, respectively). The following data are presented as mean (± standard error) mass specific corrected values. Mean AS at 10 and 15 □ was 396.30 ± 18.00 mg O^2^ kg^-1^ h^-1^ and 387.54 ± 12.76 mg O^2^ kg^-1^ h^-1^, respectively. At 23 □, AS was significantly lower than any other treatment, 138.93 ± 42.70 mg O^2^ kg^-1^ h^-1^. Mean mass specific MMR was 477.71 ± 17.19 mg O^2^ kg^-1^ h^-1^ at 10 °C and 498.66 ± 13.01 mg O^2^ kg^-1^ h^-1^ at 15 °C. Contrarily, SMR was found to increase with acclimation temperature (Figure 3-B), with mean SMR at 5 °C found to be 31.022 ± 5.034 mg O^2^ kg^-1^ h^-1^ and at its highest at 23 °C, 174.23 ± 25.64 mg O^2^ kg^-1^ h^-1^.

**Figure 3.**
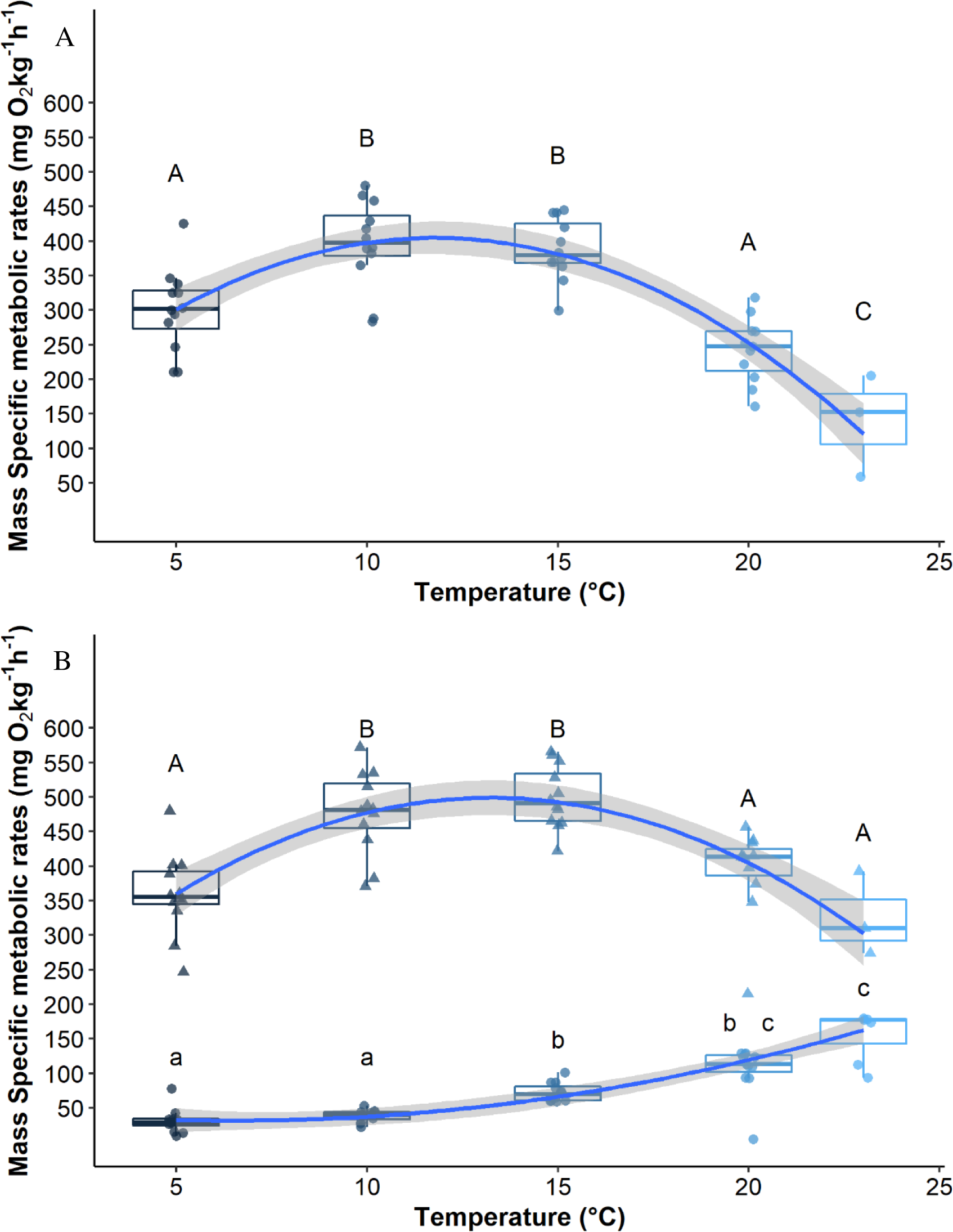
Mass specific metabolic rate estimates (Plot A Aerobic Scope, Plot B Standard Metabolic Rate and Maximum Metabolic Rate) for juvenile Brook Trout across an ecologically relevant range of acclimation temperatures. Temperatures that share a letter are not significantly different. In Plot B triangles and uppercase letters represent maximum metabolic rate and circles and lower case letters represent standard metabolic rate. Boxplots show the median, 25th and 75th percentile values, with whiskers extending up to 1.5⋅IQR. Blue lines are LOESS smoothed regression lines to emphasize the general pattern.

### Temperature Preference

Mean temperature preference of juvenile Brook Trout (mean mass 50.97 ±14.0g) estimated with Shuttle box experiments was 15.10 ± 1.13 °C. No difference in temperature preference between day (∼7:30–18:30 h) and night were found (day:15.09 ± 1.11 °C and night: 15.11 ± 1.17 °C, respectively, *p* = 0.48).

### CTmax

*CT_max_* of juvenile Brook Trout held at 10 °C was 28.2 ± 0.4 °C. The mean ± standard deviation agitation temperature of fish tested in the agitation, B_2_ and *CT_max_* treatments was 22.0 ± 1.4 °C and the mean agitation-*CT_max_*window 6.2 °C.

### Cellular level responses

#### Plasma Lactate

The linear model of lactate concentration dependent sampling time, Fulton’s condition factor, sex and the interaction of experimental treatment and recovery time was significant (*F* = 19.77, *p* < 0.001, adjusted *R*^2^ = 0.70; Figure 4). There was reasonable normality among model residuals, but variance tended to be lower at lower fitted values, and there was high collinearity between sampling time, treatment, and recovery time. Nevertheless, experimental treatment had the greatest effect size on lactate concentration (η squared = 0.43), while the interaction of treatment and recovery time (η squared = 0.19) and recovery time on its own (η squared = 0.053) were each smaller in effect size (Supplementary Table 2). Sampling time was not significant (*p* = 0.48) and its effect size was low (η squared = 0.001). Lactate concentrations were not significantly different from baseline fish at any experimental treatment except at agitation-*CT_max_*and *CT_max_* (*p* = 0.05 each), and the T^1^ was associated with higher lactate concentrations than the T^0^ (*p* < 0.001) (Supplementary Figure 1). None of sampling time, Fulton’s condition factor, or sex were significant in the ANOVA effect size statistics (*p* > 0.05 for each variable). 95% confidence intervals showed higher plasma lactate in the *CT_max_* T^1^ group relative to all other groups, including the *CT_max_* T^0^ group (Figure 4). In every other experimental treatment in the analysis, there was no significant difference between the T^0^ and T^1^ groups.

**Figure 4.**
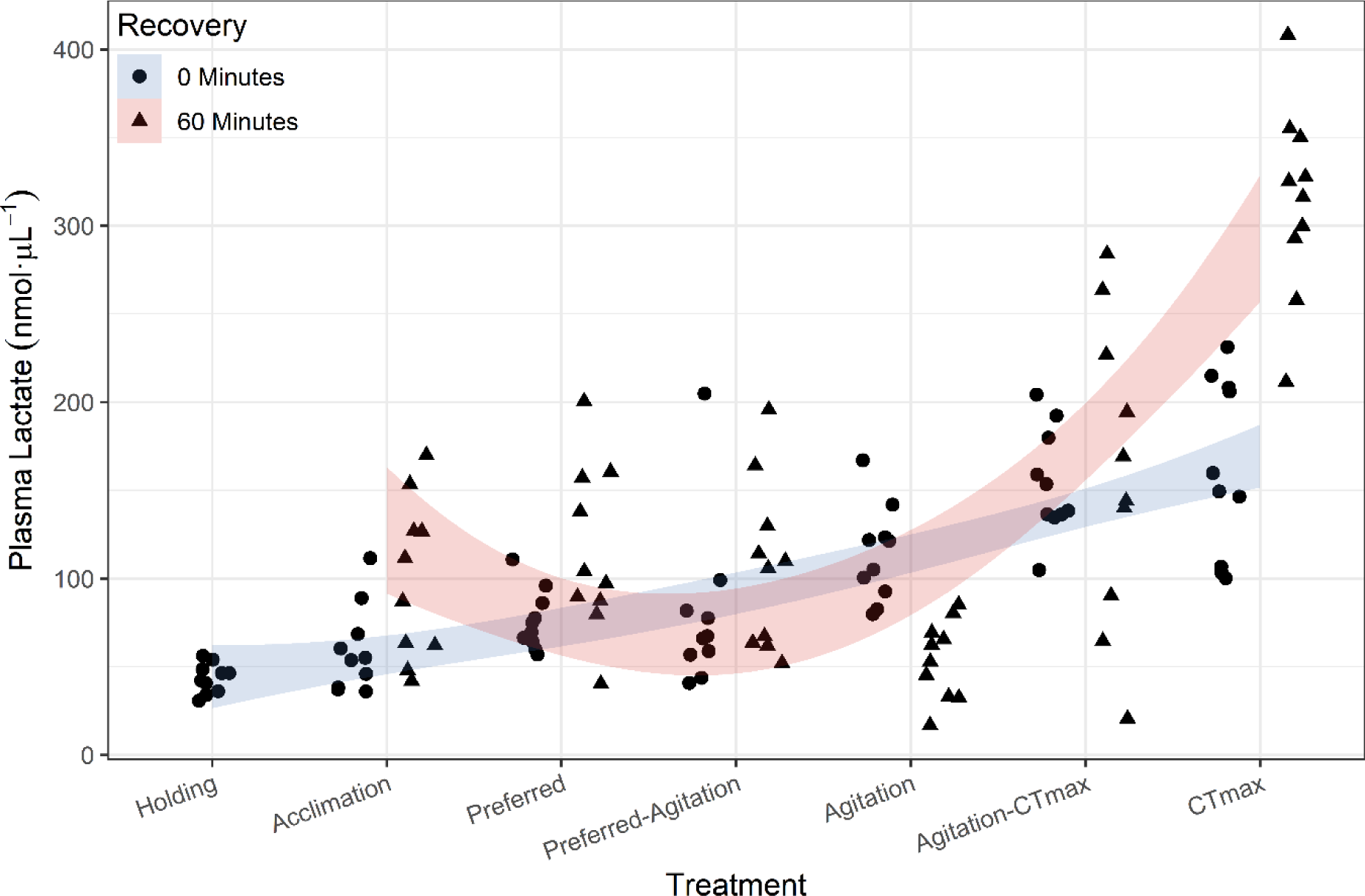
Plasma Lactate levels (nmol·μL^-1^) increased with treatment temperature and were generally higher in the 60-min recovery fish (T^1^) than the fish sampled immediately at treatment temperature (T^0^), except at the agitation temperature. Blue and red lines are LOESS smoothed regression lines for T^0^ and T^1^ groups, respectively, to emphasize the general pattern.

#### qPCR Data

In the model of mRNA abundance in gill tissue, *fos*, *hsp70a*, *ier2*, *jun*, *junb*, and *jund* showed pronounced changes both between the T^0^ and T^1^ groups and as the experiment progressed from acclimation to *CT_max_*. In the model of mRNA abundance in liver tissue, *fos*, *hsp70a*, and *ier2* showed similar pronounced changes between the groups and as the experiment progressed toward *CT_max_*.

Transcript abundance changed in both liver and gill for *hsp70a*, *ier2*, and *fos* (Figure 5). In *ier2* and *fos*, responses between the T^0^ and T^1^ groups were more substantial in the gill than the liver. In gill tissue, *hsp70a* showed a marked increase in abundance between the agitation-*CT_max_* T^0^ and T^1^ groups. This increase was consistent with a threshold effect that was both prior to *CT_max_* and subsequent to a recovery period following the agitation-*CT_max_* temperature. A similar response was observed at the agitation temperature, where in the gill, *hsp70a* and *fos* increased at agitation but only after the 60-min recovery. Therefore, we observed a transcriptional response at the same temperature as the behavioural response of agitation.

**Figure 5.**
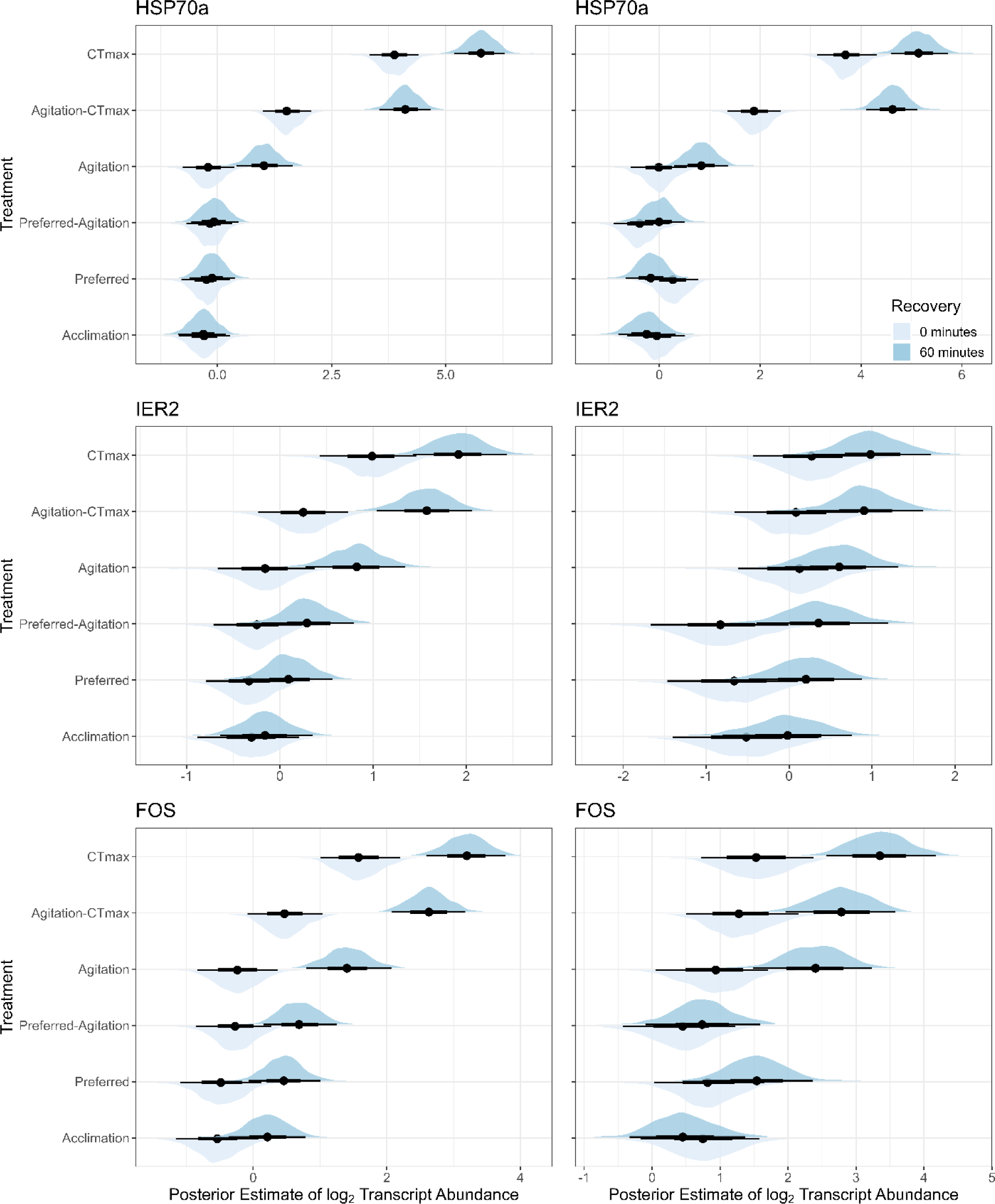
Posterior estimates of abundance of *HSP70a, IER2* and *FOS* in the gill (left) and liver (right). Transcript abundance of these genes was observed to be different at agitation temperature and higher between the 0-min (T^0^) sampling and 60-min (T^1^) sampling periods.

Transcript abundance of *jun*, *junb* and *jund* in gill tissue showed an increasing gradient in the T^1^ recovery group in all three genes (Figure 6). The T^0^groups showed no distinguishable response except in *jun* in the gill tissue between the *CT_max_* and acclimation recovery groups. In the liver, *jun* showed no significant response between any of the T^0^ versus T^1^ recovery groups. An analysis of sampling time revealed a limited association with mRNA abundance (Supplemental Results).

**Figure 6.**
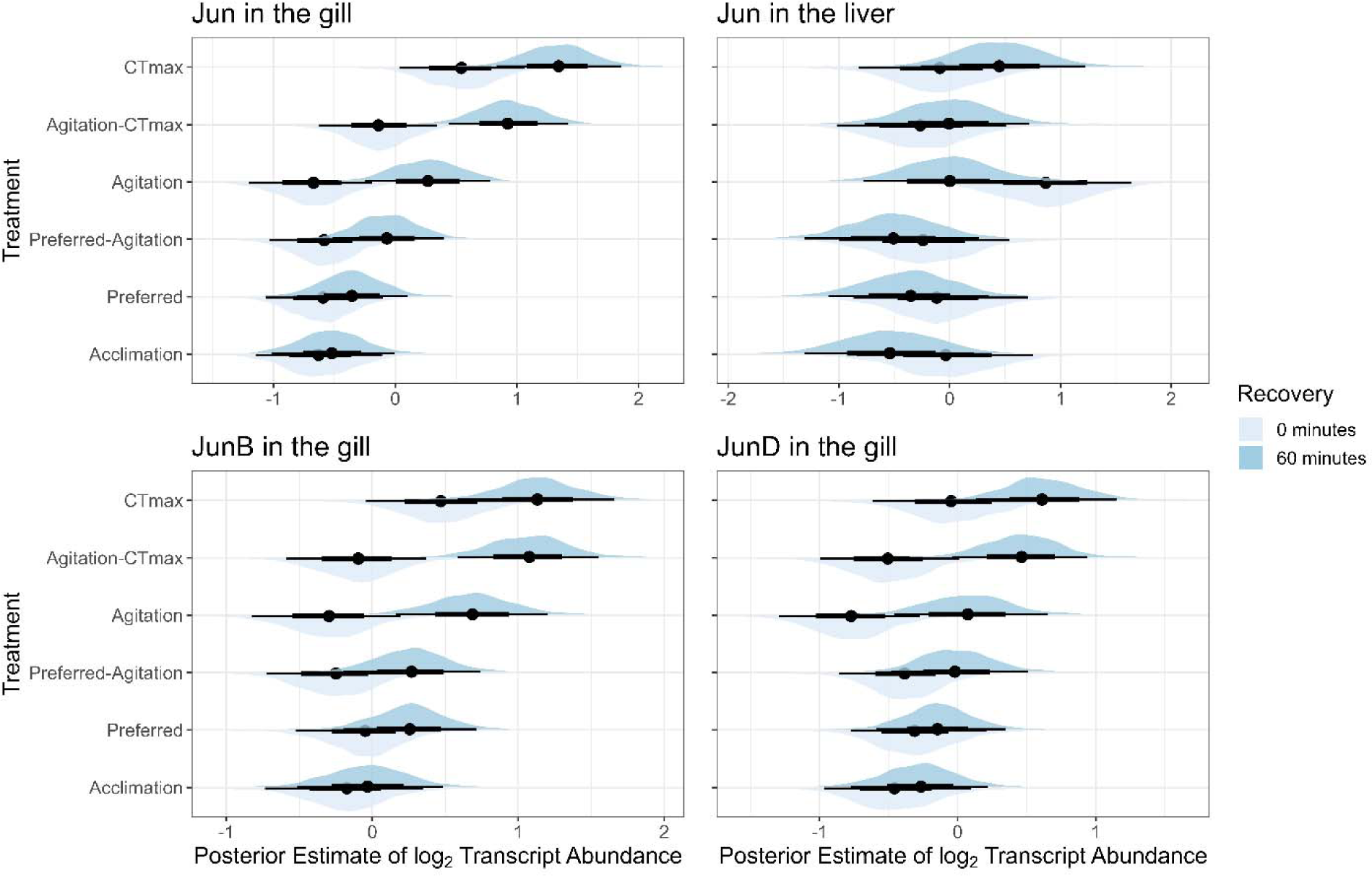
*Jun* transcript abundance in the gill tissue of juvenile Brook Trout. Lower ellipses represent tissues sampled at the 0-min (T^0^) sampling point and upper ellipses represent the 60-min (T^1^) sampling point.

## Discussion

The data from this study suggest that the agitation temperature is a useful endpoint for estimating when temperatures may start causing damage to the fish. In the current study, juvenile Brook Trout exhibited a behavioural response to temperature, declining AS, increases in plasma lactate levels and increases in stress-related mRNA transcript abundance when acutely exposed to temperatures around 22 °C. The estimated realized thermal niche of Brook Trout in the wild is predicted to be 10–20 °C (reviewed in Smith & Ridgway, 2019) and they are thought to be restricted by an upper temperature limit of ∼24 °C in nature (MacCrimmon & Campbell, 1969; Meisner, 1990; Ricker, 1934). Studies that estimated the onset of thermal avoidance and physiological stress in Brook Trout estimate an upper temperature threshold of 21–23.5 °C (Chadwick et al., 2015; Chadwick & McCormick, 2017; Goyer et al., 2014; Lund et al., 2003; reviewed in Smith and Ridgeway, 2019). The agitation temperature of 22.0 ± 1.4 °C from this study falls within this expected range, and the activation of several genes at the agitation treatment temperature, suggests that a shift in the cellular response to temperature occurs at the onset of thermal avoidance behaviour. Similar results were observed in Pacific spiny dogfish (*Squalus suckleyi*) where they showed a cellular stress response occurred at the agitation temperature (Bouyoucos et al., 2023). Morrison et al. (2020) showed that *CT_max_* generally increased with acclimation temperature in Brook Trout, however, *CT_max_* plateaued at acclimation temperatures above 20 °C that coincided with a decrease in hepatosomatic index, and an increase in plasma lactate. The results by Morrison et al. (2020) agree with the 22 °C agitation temperature and plasma lactate responses from this study despite the current study being at an acute level of exposure. One caveat on the differences seen across treatments for the changes in the mRNA transcripts in this study, is that the higher a temperature of a treatment, the longer a fish was exposed to increasing water temperatures. Therefore, there is a chance some of the responses seen may be a result of longer acute exposure to elevated temperature in the extreme temperature treatments.

A neuroendocrine response is a likely precursor to the onset of behavioural and cellular responses to high temperatures observed in this study. A neuroendocrine stress response may lead to direct tissue stimulation by the nervous system or to an increase in catecholamines released into general circulation when a stress threshold is hit, which may be followed by a glucocorticoid response (reviewed by Fabbri et al., 1998; Molinoff & Axelrod, 1971; Reid et al., 1998). A Rainbow Trout (*Onchorhynchus mykiss*) strain selected for a high cortisol response showed an increase in circulating plasma catecholamines after an acute thermal stress that was approximately 3–4 °C lower than the *CT_max_* for that strain (Leblanc et al., 2012). An increase in circulating levels of catecholamines at a temperature threshold may lead to increases in heart rate that might be expected to be associated with increased activity in the fish. Indeed, intraperitoneal injections of pharmacological agents to stimulate adrenergic receptors results in an increase in heart rate in coho salmon (*Oncorhynchus kisutch*; Casselman et al., 2012). Further, the point during acute temperature increases leading to cardiac arrhythmia occurs approximately 2–3 °C lower than the *CT_max_* in Arctic Char (*Salvelinus alpinus*; Gilbert & Farrell, 2021). These studies suggest that the effects of temperature on heart rate is under neuroendocrine control, which may be consistent with the onset of avoidance behaviours at the agitation temperature. The onset of the agitation response may also be linked to the start of neuronal dysfunction in the fish, where a behavioural avoidance response may help avoid neural impairment of the locomotor functions at higher water temperatures (Andreassen et al., 2022). Neuron activity related to increasing water temperatures has been described in other fish species (i.e., Transient Receptor Potential cation channel (TRPV1) in Zebrafish (*Danio rerio*); Gau et al., 2013), which may also be related to an activation of neuroendocrine response that is a precursor to an avoidance response at a relevant thermal limit (i.e., the agitation temperature). These neuronal responses to water temperature may be a mechanistic link between the behavioural and cellular level responses seen in this study.

We observed a delay in the cellular response in the agitation treatment where there was significantly altered mRNA transcript levels for *hsp70a* and stress inducible transcription factors after 60 min of recovery, but not at 0 min when the temperature was reached. This is in contrast to the higher temperature treatments that showed altered mRNA transcript levels when the temperature was reached (i.e., 0 min) and after 60 min of recovery. The observation of a delay in the cellular response may be consistent with the activation of a neuroendocrine response around the agitation temperature for these fish. If there was an increase in circulating cortisol, there may be a delay in when it binds to glucocorticoid receptors in target tissues leading to a cellular response that is staggered from the more rapid catecholamine response. This is consistent with previous work on pink salmon (*Oncorhynchus gorbuscha*) and sockeye salmon (*Oncorhynchus nerka*) that showed a peak in mRNA transcript abundance occurring after the peak in blood circulating cortisol after a handling stressor (Donaldson et al., 2014). Interestingly, the cortisol response has been shown to dampen the HSP response in fishes (Basu et al., 2001). A delay in the HSP70 production is consistent with a peak in circulating cortisol levels in rainbow trout when temperatures reached 25 °C (3–4 °C lower than the *CT_max_*) during acute warming from 13 °C over 1.5 h, however, peak HSP70 protein levels in the red blood cells were detected after 8 h of recovery (LeBlanc et al. 2012). Collectively, these results and the results from the present study suggest that there may be a delay between the initiation of a neuroendocrine response and peak cellular transcriptome response during an acute temperature stress, which is consistent with the cellular response being detected only after 60 min of recovery after the Brook Trout were acutely exposed to their agitation temperature.

An increase in plasma lactate turnover rates at a sub-lethal temperature threshold may explain the decrease in plasma lactate observed in the agitation temperature 60 min samples. Mackey et al. (2021) observed a 70% decrease in white muscle lactate below baseline levels in Brook Trout 24 h following exhaustive exercise at 23 °C, suggesting an increase in lactate clearance at 23 °C. Results from this study were similar, with plasma lactate levels in the 22 °C 60 min recovery treatment below baseline treatment values. Optimal clearance of plasma lactate may occur around 22 °C (i.e., agitation temperature) in this study due to a combination of an increase in plasma catecholamine related to the thermal stress response (reviewed by Fabbri et al., 1998; Molinoff & Axelrod, 1971; Reid et al., 1998), along with *in situ* glycogenesis in the white muscle, which accounts for 80–85% of the lactate produced (Milligan, 1996; reviewed by Warren & Jackson, 2008). Above the agitation temperature, lactate production due to tissue oxygen demands (Gilbert & Farrell, 2021; Leblanc et al., 2012) may exceed clearance rates, causing an accumulation within the blood and muscle tissue.

The agitation temperature appears to be a useful threshold for assessing thermal ecological limits, however, other physiological factors will likely impact fish populations over chronic time scales before this temperature is reached. Chadwick and McCormick (2017) observed that growth rates begin decreasing above 16 °C and negative growth at temperatures ∼23.4 °C. Interestingly, Chadwick and McCormick (2017) also found 22 °C to coincide with drastic increases in plasma cortisol and HSP70 levels in the gill, 12- and 11-fold higher than at 16 °C, respectively. Mackey et al. (2021) also detected physiological responses such as increased plasma cortisol, glucose, and muscle lactate levels in fish exposed to temperatures above 20 °C. The Brook Trout used in this study appeared to have a relatively large agitation-*CT_max_*window (6.2 °C), which could afford them time to seek thermal refuge from extreme temperatures, unlike other salmonids such as the Westslope Cutthroat Trout (*Oncorhynchus clarkii lewisi*), whose agitation-*CT_max_* window is smaller (1.8 °C; Enders & Durhack, 2022). The activation of a neuroendocrine and behavioural response, combined with a major shift in the transcriptome response occurred near the agitation temperature, and may be a potential link between the cellular stress response and the behavioural avoidance response in the fish.

## Acknowledgements

We would like to thank Kerry Wautier for help with animal care and fish holding.

## Competing interests

None.

## AI and AI technology

None.

## Funding

This work was supported by funding from Fisheries and Oceans Canada Strategic Program for Ecosystem Research funding to ECE, a Genome Canada Large-Scale Applied Research Project grant to CA and KMJ, and an National Science and Engineering Research Council of Canada (NSERC) Discovery Grant to KMJ (05479).

## Authors’ contributions

– T.C.D. : Conceptualization; Data curation; Formal analysis; Investigation; Methodology; Project administration; Resources; Validation; Visualization; Writing – original draft; Writing – review & editing
– M.J.T : Data curation; Formal analysis; Investigation; Methodology; Validation; Visualization; Writing – original draft; Writing – review & editing
– T.E.M. : Investigation; Methodology; Writing – original draft; Writing – review & editing
– M.A. : Investigation, Writing – review & editing
– M.J.L. : Investigation; Methodology; Writing – review & editing
– C.A. : Methodology; Writing – review & editing
– E.C.E. : Conceptualization; Funding acquisition; Project administration; Resources; Supervision; Writing – review & editing
– K.M.J. : Conceptualization; Funding acquisition; Methodology; Project administration; Resources; Supervision; Validation; Writing – original draft; Writing – review & editing

## Data Availability

Data can be made available upon request.

## Notes

### Competing Interest Statement

The authors have declared no competing interest.

## References

Andreassen, A. H., Hall, P., Khatibzadeh, P., Jutfelt, F., & Kermen, F. (2022). Brain dysfunction during warming is linked to oxygen limitation in larval zebrafish. Proceedings of the National Academy of Sciences of the United States of America, 119(39), 1–10. 10.1073/pnas.2207052119

Anttila, K., Casselman, M. T., Schulte, P. M., & Farrell, A. P. (2013). Optimum Temperature in Juvenile Salmonids: Connecting Subcellular Indicators to Tissue Function and Whole-Organism Thermal Optimum. Physiological and Biochemical Zoology, 86(2), 245–256. 10.1086/669265

Barton, K. (2020). MuMIn: Multi-Model Inference (1.43.17). https://cran.r-project.org/web/packages/MuMIn/MuMIn.pdf

Basu, N., Nakano, T., Grau, E. G., & Iwama, G. K. (2001). The Effects of Cortisol on Heat Shock Protein 70 Levels in Two Fish Species. General and Comparative Endocrinology, 124(1), 97–105. 10.1006/gcen.2001.7688

Beamish, F. W. H., & Mookherjii, P. S. (1964). Respiration of Fishes with Special Emphasis on Standard Oxygen Consumption: I. Influence of Weight and Temperature on Respiration of Goldfish, Carassius Auratus L. Canadian Journal of Zoology, 42(2), 161–175. 10.1139/z06-901

Beitinger, T., Bennett, W., & McCauley, R. (2000). Temperature tolerances of North American freshwater fishes exposed to dynamic changes in temperature. Environmental Biology of Fishes, 58, 237–275. 10.1023/A:1007676325825

Bouyoucos, I. A., Weinrauch, A. M., Jeffries, K. M., & Anderson, W. G. (2023). Physiological responses to acute warming at the agitation temperature in a temperate shark. Journal of Experimental Biology. 10.1242/jeb.246304

Bowerman, T. E., Keefer, M. L., & Caudill, C. C. (2021). Elevated stream temperature, origin, and individual size influence Chinook salmon prespawn mortality across the Columbia River Basin. Fisheries Research, 237(January), 105874. 10.1016/j.fishres.2021.105874

Brett, J. R., & Groves, T. D. D. (1979). Physiological energetics. In Fish Physiology (pp. 280–352).

Buckley, B. A., Gracey, A. Y., & Somero, G. N. (2006). The cellular response to heat stress in the goby Gillichthys mirabilis: A cDNA microarray and protein-level analysis. Journal of Experimental Biology, 209(14), 2660–2677. 10.1242/jeb.02292

Cairns, M. A., Ebersole, J. L., Baker, J. P., Wigington, P. J., Lavigne, H. R., & Davis, S. M. (2005). Influence of Summer Stream Temperatures on Black Spot Infestation of Juvenile Coho Salmon in the Oregon Coast Range. Transactions of the American Fisheries Society, 134(6), 1471–1479. 10.1577/t04-151.1

Casselman, M. T., Anttila, K., & Farrell, A. P. (2012). Using maximum heart rate as a rapid screening tool to determine optimum temperature for aerobic scope in Pacific salmon Oncorhynchus spp. Journal of Fish Biology, 80(2), 358–377. 10.1111/j.1095-8649.2011.03182.x

Chabot, D., Steffensen, J. F., & Farrell, A. P. (2016). The determination of standard metabolic rate in fishes. Journal of Fish Biology, 88(1), 81–121. 10.1111/jfb.12845

Chadwick, J. G., & McCormick, S. D. (2017). Upper thermal limits of growth in brook trout and their relationship to stress physiology. Journal of Experimental Biology, 220(21), 3976– 3987. 10.1242/jeb.161224

Chadwick, J. G., Nislow, K. H., & McCormick, S. D. (2015). Thermal onset of cellular and endocrine stress responses correspond to ecological limits in brook trout, an iconic cold-water fish. Conservation Physiology, 3(1), 1–12. 10.1093/conphys/cov017

Clark, T. D., Sandblom, E., & Jutfelt, F. (2013). Aerobic scope measurements of fishes in an era of climate change: respirometry, relevance and recommendations. Journal of Experimental Biology, 216(15), 2771–2782. 10.1242/jeb.084251

Crawshaw, L. I. (1977). Physiological and Behavioral Reactions of Fishes to Temperature Change. Journal of Fisheries Research Board of Canada, 34, 730–734. http://www.nrcresearchpress.com/doi/pdf/10.1139/f77-113

Crawshaw, L. I. (1984). Low-temperature dormancy in fish. American Journal of Physiology - Regulatory Integrative and Comparative Physiology, 15(4), 479–486. 10.1152/ajpregu.1984.246.4.r479

Desforges, J. E., Birnie□Gauvin, K., Jutfelt, F., Gilmour, K. M., Eliason, E. J., Dressler, T. L., McKenzie, D. J., Bates, A. E., Lawrence, M. J., Fangue, N., & Cooke, S. J. (2023). The Ecological Relevance of Critical Thermal Maxima Methodology (CTM) for Fishes. Journal of Fish Biology, February, 1–17. 10.1111/jfb.15368

Donaldson, M. R., Hinch, S. G., Jeffries, K. M., Patterson, D. A., Cooke, S. J., Farrell, A. P., & Miller, K. M. (2014). Comparative Biochemistry and Physiology, Part A Species- and sex-specific responses and recovery of wild, mature pacific salmon to an exhaustive exercise and air exposure stressor. *Comparative Biochemistry and Physiology*, Part A, 173, 7–16. 10.1016/j.cbpa.2014.02.019

Durhack, T. C., Mochnacz, N. J., Macnaughton, C. J., Enders, E. C., & Treberg, J. R. (2021). Life through a wider scope: Brook Trout (Salvelinus fontinalis) exhibit similar aerobic scope across a broad temperature range. Journal of Thermal Biology, 99(February), 102929. 10.1016/j.jtherbio.2021.102929

Enders, E. C., & Durhack, T. C. (2022). Metabolic rate and critical thermal maximum CTmax estimates for westslope cutthroat trout, Oncorhynchus clarkii lewisi. Conservation Physiology, 10(1), 1–9. 10.1093/conphys/coac071

Fabbri, E., Capuzzo, A., & Moon, T. W. (1998). The role of circulating catecholamines in the regulation of fish metabolism□: An overview. Comparative Biochemistry and Physiology Part C: Pharmacology, Toxicology and Endocrinology, 120(2), 177–192.

Fox, J., & Weisberg, S. (2019). An R Companion to Applied Regression (3rd). https://socialsciences.mcmaster.ca/jfox/Books/Companion/

Fry, F. E. J. (1947). Effects of the Environment on Animal Activity. Publications of the Ontario Fisheries Research Laboratory, 55(LXVIII), 1–62.

Fry, F. E. J. (1971). The effect of environmental factors on the physiology of fish. In W. Hoar & D. Randell (Eds.), Fish Physiology: Environmental relations and behavior (pp. 1–98). Academic Press.

Gau, P., Poon, J., Ufret-Vincenty, C., Snelson, C. D., Gordon, S. E., Raible, D. W., & Dhaka, A. (2013). The zebrafish ortholog of TRPV1 is required for heat-induced locomotion. Annals of Internal Medicine, 158(6), 5249–5260. 10.1523/JNEUROSCI.5403-12.2013

Gilbert, M. J. H., & Farrell, A. P. (2021). The thermal acclimation potential of maximum heart rate and cardiac heat tolerance in arctic char (Salvelinus alpinus), a northern cold-water specialist. Journal of Thermal Biology, 95(October 2020), 102816. 10.1016/j.jtherbio.2020.102816

Goyer, K., Bertolo, A., Pépino, M., & Magnan, P. (2014). Effects of lake warming on behavioural thermoregulatory tactics in a cold-water stenothermic fish. PLoS ONE, 9(3). 10.1371/journal.pone.0092514

Hadfield, J. D. (2010). MCMCglmm: MCMC Methods for Multi-Response GLMMs in R. Journal of Statistical Software, 33(2), 1–22. https://www.jstatsoft.org/v33/i02/

Hasnain, S. (2012). Factors influencing ecological metrics of thermal response in North American freshwater fish.

Hothorn, T., Bretz, F., & Westfall, P. (2008). Simultaneous Inference in General Parametric Models. Biometrical Journal, 50(3), 346–363.

J, L. (2006). Plotrix: a package in the red light district of R. R-News, 6(4), 8–12.

Jeffries, K. M., Connon, R. E., Davis, B. E., Komoroske, L. M., Britton, M. T., Sommer, T., Todgham, A. E., & Fangue, N. A. (2016). Effects of high temperatures on threatened estuarine fishes during periods of extreme drought. Journal of Experimental Biology, 219(11), 1705–1716. 10.1242/jeb.134528

Jeffries, K. M., Fangue, N. A., & Connon, R. E. (2018). Multiple sub-lethal thresholds for cellular responses to thermal stressors in an estuarine fish. Comparative Biochemistry and Physiology -Part A_J: Molecular and Integrative Physiology, 225(June), 33–45. 10.1016/j.cbpa.2018.06.020

Jourdain-Bonneau, C., Deslauriers, D., Gourtay, C., Jeffries, K. M., & Audet, C. (2023). Metabolic and transcriptomic response of two juvenile anadromous brook charr (. Canadian Journal of Zoology.

Kay, M. (2021). tidybayes: Tidy data and geoms for bayesian models (3.0.2). 10.5281/zenodo.1308151

Kelly, N. I., Burness, G., McDermid, J. L., & Wilson, C. C. (2014). Ice age fish in a warming world: Minimal variation in thermal acclimation capacity among lake trout (Salvelinus namaycush) populations. Conservation Physiology, 2(1), 1–14. 10.1093/conphys/cou025

Killen, S. S., Marras, S., Metcalfe, N. B., McKenzie, D. J., & Domenici, P. (2013). Environmental stressors alter relationships between physiology and behaviour. Trends in Ecology and Evolution, 28(11), 651–658. 10.1016/j.tree.2013.05.005

Komoroske, L. M., Connon, R. E., Jeffries, K. M., & Fangue, N. A. (2015). Linking transcriptional responses to organismal tolerance reveals mechanisms of thermal sensitivity in a mesothermal endangered fish. Molecular Ecology, 24(19), 4960–4981. 10.1111/mec.13373

Leblanc, S., Höglund, E., Gilmour, K. M., & Currie, S. (2012). Hormonal modulation of the heat shock response□: insights from fish with divergent cortisol stress responses. American Journal of Physiology - Regulatory Integrative and Comparative Physiology, 302(1), R184– R192. 10.1152/ajpregu.00196.2011

Life Technologies. (2011). RNAlater Tissue Collection: RNA Stabilization Solution. https://tools.thermofisher.com/content/sfs/manuals/cms_056069.pdf

Logan, C. A., & Somero, G. N. (2011). Effects of thermal acclimation on transcriptional responses to acute heat stress in the eurythermal fish Gillichthys mirabilis (Cooper). American Journal of Physiology - Regulatory Integrative and Comparative Physiology, 300(6), 1373–1383. 10.1152/ajpregu.00689.2010

Lüdecke, D. (2018). sjstats: Statistical functions for regression models (0.18.1). 10.5281/zenodo.1284472

Lüdecke, D. (2021). sjPlot: Data visualization for statistics in social science (2.8.10). https://cran.r-project.org/package=sjPlot

Lüdecke, D., Ben-Shachar, M. S., Indrajeet, P., Waggoner, P., & Makowski, D. (2021). performance: An R package for assessment, comparison and testing statistical models. The Journal of Open Source Software, 6(60), 3139. 10.21105/joss.03139

Lund, E., Olsen, E. M., & Vøllestad, L. A. (2003). First-year survival of brown trout in three Norwegian streams. Journal of Fish Biology, 62(2), 323–340. 10.1046/j.1095-8649.2003.00025.x

MacCrimmon, H. R., & Campbell, S. J. (1969). World Distribution of Brook Trout, Salaelinus fontinalis. Journal of Fisheries Research Board of Canada, 26(7), 1699–1725.

Mackey, T. E., Hasler, C. T., Durhack, T. C., Jeffrey, J. D., Macnaughton, C. J., Ta, K., Enders, E. C., & Jeffries, K. M. (2021). Molecular and physiological responses predict acclimation limits in juvenile brook trout (Salvelinus fontinalis). Journal of Experimental Biology, 224(16). 10.1242/jeb.241885

Macnaughton, C. J., Durhack, T. C., Mochnacz, N. J., & Enders, E. C. (2021). Metabolic performance and thermal preference of westslope cutthroat trout (Oncorhynchus clarkii lewisi) and non-native trout across an ecologically relevant range of temperatures. Canadian Journal of Fisheries and Aquatic Sciences, 78, 1247–1256. 10.1139/cjfas-2020-0173

Macnaughton, C. J., Kovachik, C., Charles, C., & Enders, E. C. (2018). Using the shuttlebox experimental design to determine temperature preference for juvenile Westslope Cutthroat Trout (Oncorhynchus clarkii lewisi). Conservation Physiology, 6(1), 1–10. 10.1093/conphys/coy018

Matz, M. V., Wright, R. M., & Scott, J. G. (2013). No control genes required: Bayesian analysis of qRT-PCR data. PloS One, 8(8), 1–12. 10.1371/journal.pone.0071448

Mauger, S., Shaftel, R., Leppi, J. C., & Rinella, D. J. (2017). Summer temperature regimes in southcentral Alaska streams: Watershed drivers of variation and potential implications for Pacific salmon. Canadian Journal of Fisheries and Aquatic Sciences, 74(5), 702–715. 10.1139/cjfas-2016-0076

McDonnell, L. H., & Chapman, L. J. (2015). At the edge of the thermal window: Effects of elevated temperature on the resting metabolism, hypoxia tolerance and upper critical thermal limit of a widespread African cichlid. Conservation Physiology, 3(1), 1–13. 10.1093/conphys/cov050

Meisner, J. D. (1990). Effect of Climatic Warming on the Southern Margins of the Native Range of Brook Trout, Salvelinus fontinalis. Canadian Journal of Fisheries & Aquatic Sciences, 47, 1065–1070.

Milligan, C. L. (1996). Metabolic Recovery from Exhaustive Exercise in Rainbow Trout. Comparative Biochemistry and Physiology Part A: Physiology, 113(1), 51–60. 10.1016/0300-9629(95)02060-8

Molinoff, P. B., & Axelrod, J. (1971). Biochemistry of catecholamines. Annual Review of Biochemistry, 40(1), 465–500.

Morash, A. J., Speers□Roesch, B., Andrew, S., & Currie, S. (2020). The physiological ups and downs of thermal variability in temperate freshwater ecosystems. Journal of Fish Biology. 10.1111/jfb.14655

Morrison, S. M., Mackey, T. E., Durhack, T. C., Jeffrey, J. D., Wiens, L. M., Mochnacz, N. J., Hasler, C. T., Enders, E. C., Treberg, J. R., & Jeffries, K. M. (2020). Sub-lethal temperature thresholds indicate acclimation and physiological limits in brook trout Salvelinus fontinalis. Journal of Fish Biology, 97(2), 583–587. 10.1111/jfb.14411

Munday, P. L., Crawley, N. E., & Nilsson, G. E. (2009). Interacting effects of elevated temperature and ocean acidification on the aerobic performance of coral reef fishes. Marine Ecology Progress Series, 388, 235–242. 10.3354/meps08137

Petty, J. T., Hansbarger, J. L., Huntsman, B. M., & Mazik, P. M. (2012). Brook trout movement in response to temperature, flow, and thermal refugia within a complex Appalachian riverscape. Transactions of the American Fisheries Society, 141(4), 1060–1073. 10.1080/00028487.2012.681102

Poletto, J. B., Cocherell, D. E., Baird, S. E., Nguyen, T. X., Cabrera-stagno, V., Farrell, A. P., & Fangue, N. A. (2017). Unusual aerobic performance at high temperatures in juvenile Chinook salmon, *Oncorhynchus tshawytscha*. Conservation Physiology, 5, 1–13. 10.1093/conphys/cow067

Posit Team. (2020). RStudio: Integrated Development Environment for R (1.3.1056). Posit Software, PBC. http://posit.co/

R Core Team. (2020). R: A Language and Environment for Statistical Computing (4.0.2). R Foundation for Statistical Computing. http://www.r-project.org

Reid, S. G., Bernier, N. J., & Perry, S. F. (1998). The adrenergic stress response in fish_J: control of catecholamine storage and release. 120, 1–27.

Reidy, S. P., Nelson, J. A., Tang, Y., & Kerr, S. R. (1995). Post-exercise metabolic rate in Atlantic cod and its dependence upon the method of exhaustion. Journal of Fish Biology, 47, 377–386.

Ricker, W. E. (1934). An ecological classification of certain Ontario streams. Ontario Fisheries Research Laboratory, Publication No 49, University of Toronto Studies, Biological Series, No 37. University of Toronto Press. Toronto, ON.

Rodnick, K. J., St.-Hilaire, S., Battiprolu, P. K., Seiler, S. M., Kent, M. L., Powell, M. S., & Ebersole, J. L. (2008). Habitat Selection Influences Sex Distribution, Morphology, Tissue Biochemistry, and Parasite Load of Juvenile Coho Salmon in the West Fork Smith River, Oregon. Transactions of the American Fisheries Society, 137(6), 1571–1590. 10.1577/t07-138.1

Schreck, C. B. (2010). Stress and fish reproduction: The roles of allostasis and hormesis. General and Comparative Endocrinology, 165(3), 549–556. 10.1016/j.ygcen.2009.07.004

Schulte, P. M. (2014). What is environmental stress? Insights from fish living in a variable environment. Journal of Experimental Biology, 217, 23–34. 10.1242/jeb.089722

Schulte, P. M., Healy, T. M., & Fangue, N. A. (2011). Thermal performance curves, phenotypic plasticity, and the time scales of temperature exposure. Integrative and Comparative Biology, 51(5), 691–702. 10.1093/icb/icr097

Smith, D. A., & Ridgway, M. S. (2019). Temperature selection in Brook Charr: lab experiments, field studies, and matching the Fry curve. Hydrobiologia, 840(1), 143–156. 10.1007/s10750-018-3869-4

Tomanek, L. (2010). Variation in the heat shock response and its implication for predicting the effect of global climate change on species’ biogeographical distribution ranges and metabolic costs. Journal of Experimental Biology, 213(6), 971–979. 10.1242/jeb.038034

Treberg, J. R., Killen, S. S., MacCormack, T. J., Lamarre, S. G., & Enders, E. C. (2016). Estimates of metabolic rate and major constituents of metabolic demand in fishes under field conditions: Methods, proxies, and new perspectives. Comparative Biochemistry and Physiology -Part A_J: Molecular and Integrative Physiology, 202, 10–22. 10.1016/j.cbpa.2016.04.022

Venables, W. N., & Ripley, B. D. (2002). Modern Applied Statistics with S (4th ed.). Springer. http://www.stats.ox.ac.uk/pub/MASS4

Warren, D. E., & Jackson, D. C. (2008). Lactate metabolism in anoxic turtles□: an integrative review. Journal of Comparative Physiology B, 178, 133–148. 10.1007/s00360-007-0212-1

Wells, Z. R. R., McDonnell, L. H., Chapman, L. J., & Fraser, D. J. (2016). Limited variability in upper thermal tolerance among pure and hybrid populations of a cold-water fish. Conservation Physiology, 4(1), cow063. 10.1093/conphys/cow063

Westley, P. A. H. (2020). Documentation of en route mortality of summer chum salmon in the Koyukuk River, Alaska and its potential linkage to the heatwave of 2019. Ecology and Evolution, 10(19), 10296–10304. 10.1002/ece3.6751

White, S. L., & Wagner, T. (2021). Behaviour at short temporal scales drives dispersal dynamics and survival in a metapopulation of brook trout (Salvelinus fontinalis). Freshwater Biology, 66(2), 278–285. 10.1111/fwb.13637

Wickham, H., Averick, M., Bryan, J., Chang, W., McGowan, L., François, R., Grolemund, G., Hayes, A., Henry, L., Hester, J., Kuhn, M., Pedersen, T. L., Miller, E., Bache, S. M., Müller, K., Ooms, J., Robinson, D., Seidel, D. P., Spinu, V., … Yutani, H. (2019). Welcome to the tidyverse. Journal of Open Source Software, 4(43), 1.3.0. 10.21105/joss.01686

Wickham, H., Francois, R., Henry, L., & Muller, K. (2020). A Grammar of Data Manipulation (0.8.5). http://dplyr.tidyverse.org/

