## Supplementary Material for "Behavioural responses to acute warming precede critical shifts in the cellular and physiological thermal stress responses in fish"

Supplementary Data

**Methods**

*qPCR Data*

To assess the possibility that changes in mRNA abundance may reflect duration of the experiment as opposed to increasing temperatures, linear models were used to investigate associations between time and mRNA abundance in both gill and liver. These linear models of transcript abundance dependent on time in minutes from the start of experimental manipulations to blood draw (sampling time) were used with every gene in both gill and liver data. Here, a ‘one factor’ model where treatment and recovery conditions were combined into a single variable. Fish were haphazardly sampled from holding tanks were used as the reference treatment group against which other treatments were compared. These models were used to assess transcript abundance changes in the acclimation group, which were pulled from holding tanks into experimental tanks approximately 18 hours prior to the experiment and may have induced changes in mRNA transcript abundance.

**Results**

*qPCR Data*

Among linear models of mRNA abundance dependent on sampling time, 4 genes (out of 20) were significant in liver tissue and 6 (out of 22) were significant in gill tissue (Supplementary Tables 3 and 4). The highest adjusted *R*^2^ values were 0.26 and 0.30 for *hsp70a* in the gill and liver, respectively (gill: *F* = 62, *p* < 0.001; liver: *F* = 80, *p* < 0.001). However, correlation coefficients were negative for each model of *hsp70a* in the gill and liver (-0.51 and -0.55, respectively), indicating that increased sampling time was correlated with a decrease in *hsp70a* transcript abundance, not an increase in transcript abundance as might be predicted if experimental duration induced a stress response on its own. Only a single gene among any of the linear models showed a positive correlation between sampling time and mRNA abundance: *mycb* in gill tissue (*F* = 4.5, *p* = 0.036, adjusted *R*^2^ = 0.020) with a correlation coefficient of 0.16. Given the few genes with significant associations between mRNA abundance and sampling time, the generally negative correlations between mRNA abundance and sampling time in the significant associations that were observed, and the low values for variance explained (adjusted *R*^2^) in mRNA abundance by sampling time, we concluded that sampling time likely had a mixed but limited association with gene expression. Given its significant associations in several genes and causal potential for modifying gene expression, sampling time was nevertheless included in the Bayesian analyses of differential expression as a fixed effect.

One factor models showed that mRNA abundance of any gene in fish from holding tanks did not differ from mRNA abundance of any gene in fish of the acclimation 0-minute recovery group (Supplementary Figures 2 & 3). Therefore, we concluded that fish transfer from holding to experimental tanks ~18 hours prior to experimental did not induce transcriptional changes among the genes used in the present qPCR panel.

Supplementary Tables

Supplementary Table 1. Primer and probe sequences for quantitative PCR in brook trout (Salvelinus fontinalis).

| **Gene name** | **Gene Abbreviation** | **Primer Sequence (5’–3’)** | **Product size (bp)** | **Eff. (%)** |
| --- | --- | --- | --- | --- |
| *caspase-9* | *casp9* | F: ATGTCCTCCAGCAGTGACTCTCT  R: GGGTAGTGTGGCCTTTGCA  Probe: AGCACTCAGTCTGATGAG | 66 | 103^G^; 101^L^ |
| *catalase* | *cat* | F: TGTGCATGTTGGACAACCA  R: GGTCTCTGGAGCACTGAAGCT  Probe: CTGGCGCTCCCAACTACTTCCCCA | 68 | 106^G^; 101^L^ |
| *cold-inducible RNA-binding protein-a* | *cirbpa* | F: GGGAAGGTCTCGTGGATTCG  R: GGTTCTGCCATCGACAGACT  Probe: GGACGCATTGGAGGGAATGAACGGC | 100 | 105^G^; 106^L^ |
| *citrate synthase* | *cs* | F: TGTCCTTCAGCGCAGCTATG  R: TCCTGGTTAGCCAGTCCATGA  Probe: CGGATTGGCCGGACC | ? | 100^G^; 103^L^ |
| *elongation factor 1-alpha* | *ef1a* | F: GAAGCTTGAGGACAACCCCA  R: AAGCTCTCCACACACATGGG  Probe: CGCCGCCATCATCGTCATGGTGC | 90 | 105^G^; 103^L^ |
| *protein c-fos* | *fos* | F: GCCTGAGGAGGAGGAGAAGA  R: TGCAGAGTGTCGGTGAGTTC  Probe: GCAGCAGCTAAATGCCGCAACAGGA | 99 | 90^G^; 102^L^ |
| *glutathione peroxidase 1a* | *gpx1a* | F: CACCCCTTGTTTGTGTATCTCA  R: GGGCTCCACATGATGAACTT  Probe: ATAAATTGCCATTCCCCTCCGATGACTC | 95 | 92^G^; 92^L^ |
| *heat shock transcription factor 1* | *hsf1* | F: CCCAAGTTCAGCAGGCAGTA  R: CGTGAAGAGACCGGTACTGG  Probe: CCTCGCTGCAGGGCTCGCCT | 108 | 93^G^; 103^L^ |
| *heat shock protein 70-alpha* | *hsp70a* | F: GAACACTGTCCTCCAGCTCC  R: CCCTGAAGAGGTCGGAACAC  Probe: ACACCTCCATCACCAGGGCTCGTTT | 120 | 99^G^; 100^L^ |
| *cytosolic heat shock protein 90-alpha* | *hsp90aa* | F: ATTCCCAACAAGGAGGAGCG  R: GGTGCCAGACTTTGCAATGG  Probe: CGGCATCGGCATGACCAAGGCC | 105 | 89^L^ |
| *heat shock protein 90-alpha-beta-1* | *hsp90ab1* | F: GGCCAAGAAACACCTGGAGAT  R: TGCCTCAGGGTCTCCACAA  Probe: AACCCAGACCACCCC | 57 | 97^G^; 103^L^ |
| *heat shock protein 90-beta-alpha* | *hsp90ba* | F: AGGTGGAGGAGGACGAGTAC  R: GCTGTGAAGTGGATGTGGGA  Probe: TCTCCAGGGACACAGACGAGCCCA | 85 | 99^G^; 102^L^ |
| *immediate-early response gene 2* | *ier2* | F: CCTGCGATGCGTGGTAGATA  R: GCAAACTATACAGCTCGCGC  Probe: CCGGTGGAGACGGTGTCCCCC | 108 | 87^G^; 89^L^ |
| *transcription factor ap-1* | *jun* | F: ACCCAACCAGCACACTCAAA  R: CTGGGGAGGCCAGTTTAAGG  Probe: GCCAGCGACATCCTCACCTCCCC | 93 | 85^G^; 97^L^ |
| *transcription factor jun-b* | *junb* | F: GCCAGCGACATCCTCACCTCCCC  R: CTGCTCTCCATGTCTGGGAC  Probe: GGAGCAGGCCTACACGCACAGC | 94 | 93^G^ |
| *transcription factor jun-d* | *jund* | F: CGACATGCAGTGCTTTGGAG  R: CTTTGATGCGCTCCTGGTTG  Probe: AGCCCGCCGTTGTCTCCTATTGACA | 71 | 107^G^ |
| *lactate dehydrogenase* | *ldh* | F: AGCGTCCTCCTCAGGGACTT  R: AGCTTATCCTCCATCACGTCAAC  Probe: CTGATGAGCTGGCTCT | ? | 102^G^; 96^L^ |
| *transcriptional regulator myc-2* | *mycb* | F: TGATGAGGAGGAGGACGAGG  R: AGCACAAGGGGACTGTGATG  Probe: AGCGGTCCGACCCCAGCACG | 114 | 90^G^ |
| *60S ribosomal protein L7* | *rpl7* | F: CCATGGCAGGATGACCAA  R: CCCAGAGCCTTATCGATCAG  Probe: CAGCGTATCGCCCTCACAGACAACG | 66 | 105^G^; 100^L^ |
| *60S ribosomal protein L13a* | *rpl13a* | F: CACTGGAGAGGCTGAAGGTG  R: GTGGGCTTCAGACGGACAAT  Probe: GCGCATGGTCGTACCTGCTGCC | 103 | 97^G^; 102^L^ |
| *40S ribosomal protein S9* | *rps9* | F: TCTCCCTGCGTTCACCATAC  R: GCCCTTCTTGGCATTCTTTC  Probe: ACGCCCCGGCCGTGTCAA | 68 | 97^G^; 99^L^ |
| *superoxide dismutase [Cu-Zn]* | *sod1* | F: CAACACCAACGGCTGTATGA  R: CTCCGTGGGTCTTGTTGTG  Probe: TGCCGGACCCCACTTCAACCC | 62 | 100^G^; 102^L^ |
| *superoxide dismutase [Mn]* | *sod2* | F: ATGGCTGGGCTTTGACAA  R: CTGCAGTGGGTCTTGATTAGG  Probe: AGCGGGAAGCTCCGTATCACAGCC | 70 | 103^G^; 97^L^ |

^L^ denotes liver

^G^ denotes gill

Forward and reverse primer sequences are indicated by “F” and “R”, respectively.

Supplementary Table 2. Plasma lactate Anova statistical analysis table.

| Variable | Sum of squares | Mean squares | Degrees of freedom | F statistic | p value | eta squared |
| --- | --- | --- | --- | --- | --- | --- |
| Sampling time | 962.702 | 962.702 | 1 | 0.5 | 0.481 | 0.001 |
| Fulton’s condition factor | 2727.317 | 2727.317 | 1 | 1.418 | 0.236 | 0.004 |
| Sex | 2442.983 | 1221.491 | 2 | 0.635 | 0.532 | 0.004 |
| treatment | 279888.3 | 46648.04 | 6 | 24.249 | 0 | 0.428 |
| Recovery time | 34450.03 | 34450.03 | 1 | 17.908 | 0 | 0.053 |
| Treatment * recovery time | 116673.2 | 23334.64 | 5 | 12.13 | 0 | 0.178 |
| Residuals | 217383.2 | 1923.745 | 113 | N/A | N/A | N/A |

Supplementary Table 3. Gill gene expression linear model full statistical analysis. Bolded P values were significantly different.

| gene | p value | F statistic | Degrees of Freedom | Adjusted R^2^ | Correction coefficient |
| --- | --- | --- | --- | --- | --- |
| rpl13a | 0.453776 | 0.563773 | 1, 171 | -0.00254 | 0.057324 |
| rps9 | 0.926901 | 0.008442 | 1, 171 | -0.0058 | 0.007026 |
| rpl7 | 0.758479 | 0.094846 | 1, 171 | -0.00529 | 0.023545 |
| cirbpa | 0.212655 | 1.56492 | 1, 171 | 0.003274 | 0.095229 |
| ef1a | 0.2594 | 1.280447 | 1, 171 | 0.001628 | 0.086211 |
| hsf1 | 0.082526 | 3.050132 | 1, 171 | 0.011779 | 0.13238 |
| hsp70a | **4.36E-13** | 61.61906 | 1, 171 | 0.260594 | -0.51468 |
| hsp90ba | 0.769386 | 0.086226 | 1, 171 | -0.00534 | -0.02245 |
| casp | 0.756761 | 0.096245 | 1, 171 | -0.00528 | -0.02372 |
| cat | 0.303041 | 1.067185 | 1, 171 | 3.90E-04 | 0.078754 |
| cs | 0.599131 | 0.277342 | 1, 171 | -0.00422 | 0.04024 |
| gpx1a | 0.451053 | 0.570616 | 1, 171 | -0.0025 | 0.05767 |
| hsp90ab1 | 0.79465 | 0.067956 | 1, 171 | -0.00545 | 0.019931 |
| ldh | 0.960812 | 0.002421 | 1, 171 | -0.00583 | 0.003763 |
| sod1 | 0.883947 | 0.02137 | 1, 171 | -0.00572 | -0.01118 |
| sod2 | 0.709342 | 0.139399 | 1, 171 | -0.00503 | 0.02854 |
| jund | 0.430131 | 0.625432 | 1, 171 | -0.00218 | -0.06037 |
| mycb | **0.035604** | 4.486748 | 1, 171 | 0.019869 | 0.159898 |
| junb | **0.01321** | 6.271002 | 1, 171 | 0.029734 | -0.18808 |
| jun | **5.30E-04** | 12.4777 | 1, 171 | 0.062556 | -0.26078 |
| ier2 | **9.83E-07** | 25.7998 | 1, 171 | 0.126015 | -0.36207 |
| fos | **2.50E-07** | 28.87615 | 1, 171 | 0.139467 | -0.38009 |

Supplementary Table 4. Liver gene expression linear model full statistical analysis. Bolded P values were significantly different.

| gene | p value | F Statistic | Degrees of Freeedom | Adjusted R^2^ | Correction coefficient |
| --- | --- | --- | --- | --- | --- |
| rpl13a | 0.655467 | 0.199713 | 1, 188 | -0.00425 | -0.03258 |
| rps9 | 0.820114 | 0.051855 | 1, 188 | -0.00504 | -0.01661 |
| rpl7 | 0.728018 | 0.121301 | 1, 188 | -0.00467 | 0.025393 |
| cirbpa | 0.829981 | 0.046236 | 1, 188 | -0.00507 | 0.01568 |
| ef1a | 0.831666 | 0.04531 | 1, 188 | -0.00508 | -0.01552 |
| hsf1 | 0.796289 | 0.066832 | 1, 188 | -0.00496 | 0.018851 |
| hsp70a | **3.39E-16** | 80.11555 | 1, 188 | 0.29508 | -0.54663 |
| hsp90ba | **0.036267** | 4.447969 | 1, 188 | 0.017916 | -0.15203 |
| casp | 0.379323 | 0.776551 | 1, 188 | -0.00118 | 0.064137 |
| cat | 0.131175 | 2.29854 | 1, 188 | 0.006824 | 0.109903 |
| cs | 0.810957 | 0.057374 | 1, 188 | -0.00501 | -0.01747 |
| gpx1a | 0.156069 | 2.028118 | 1, 188 | 0.00541 | 0.103309 |
| hsp90ab1 | 0.227802 | 1.464074 | 1, 188 | 0.002449 | 0.087906 |
| ldh | 0.621402 | 0.244711 | 1, 188 | -0.00401 | -0.03605 |
| sod1 | 0.236937 | 1.40769 | 1, 188 | 0.002152 | 0.086209 |
| sod2 | 0.295704 | 1.099582 | 1, 188 | 5.27E-04 | 0.076255 |
| hsp90aa | 0.712113 | 0.136602 | 1, 188 | -0.00474 | 0.027386 |
| fos | **2.02E-06** | 24.05806 | 1, 188 | 0.10925 | -0.33762 |
| ier2 | **1.88E-04** | 14.55337 | 1, 188 | 0.070755 | -0.27564 |
| jun | 0.616479 | 0.251693 | 1, 188 | -0.00402 | -0.03676 |

Supplementary Figures

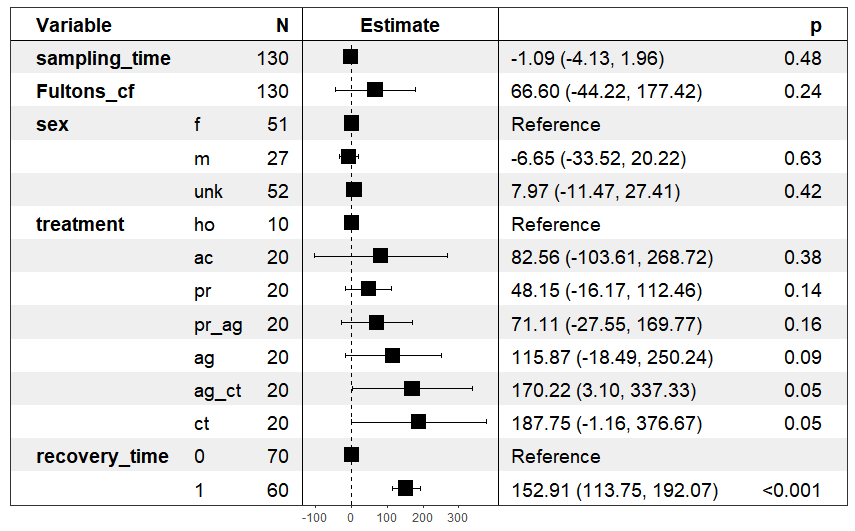

Supplementary Figure 1 – Plasma Lactate forest plot.

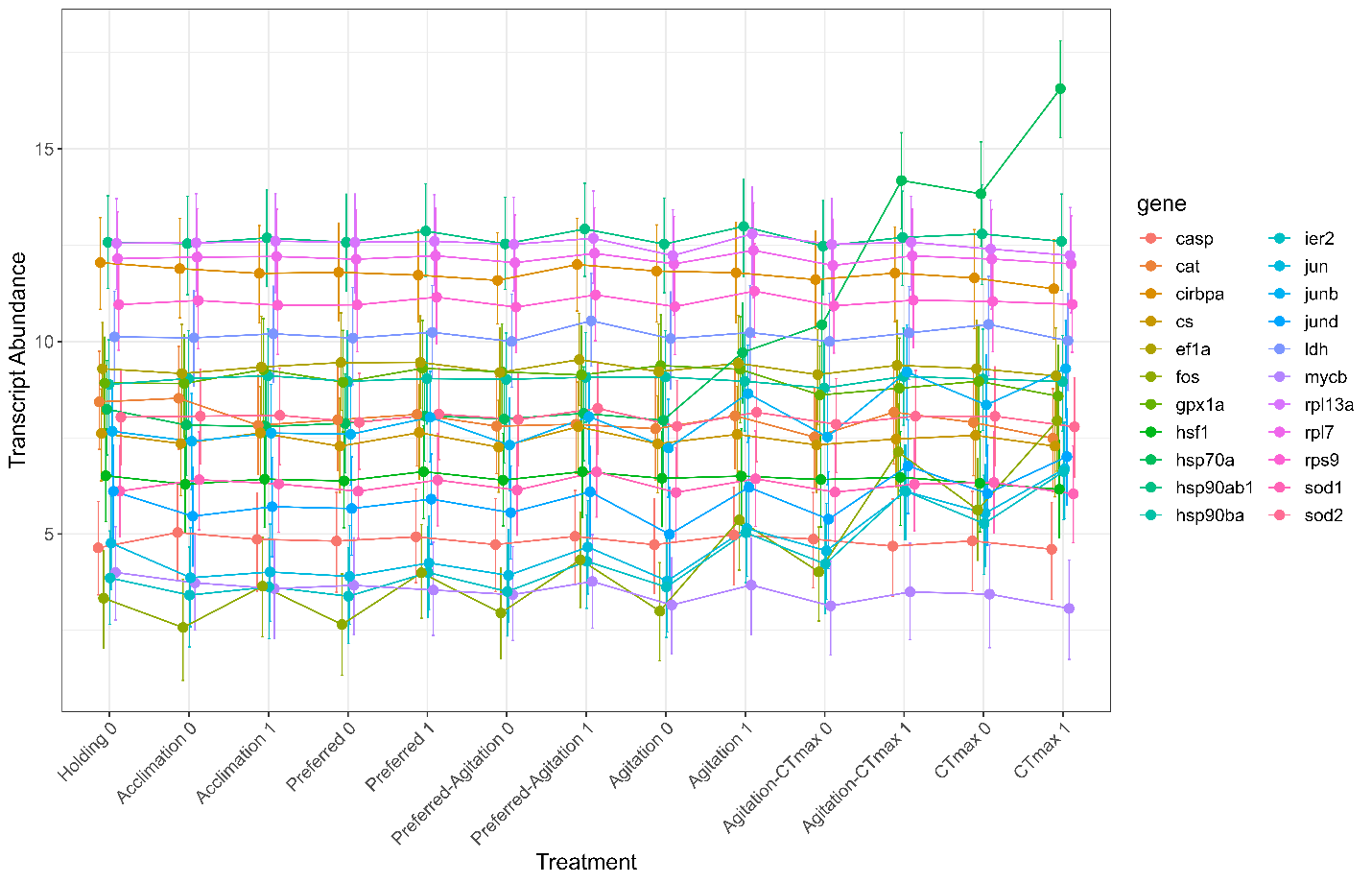

Supplementary Figure 2 – Gill transcription abundance regulation from 1-factor analysis.

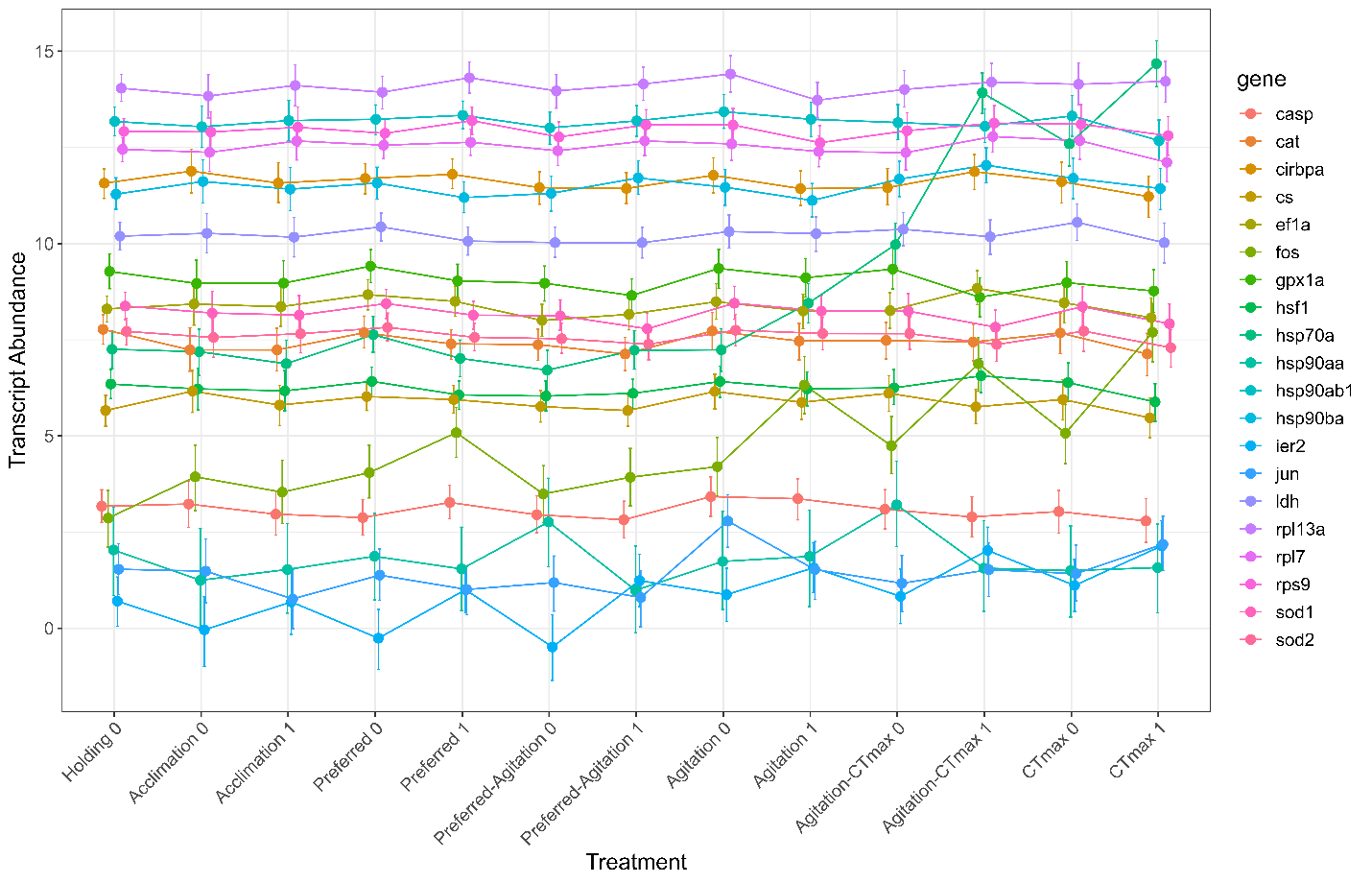

Supplementary Figure 3 – Liver transcript abundance regulation from 1-factor analysis.
